# Paradoxes and synergies: optimizing management of a deadly virus in an endangered carnivore

**DOI:** 10.1101/2021.01.11.426249

**Authors:** Marie L.J. Gilbertson, Dave Onorato, Mark Cunningham, Sue VandeWoude, Meggan E. Craft

## Abstract

1. Pathogen management strategies in wildlife are typically accompanied by an array of uncertainties such as the efficacy of vaccines or potential unintended consequences of interventions. In the context of such uncertainties, models of disease transmission can provide critical insight for optimizing pathogen management, especially for species of conservation concern. The endangered Florida panther experienced an outbreak of feline leukemia virus (FeLV) in 2002-04, and continues to be affected by this deadly virus. Ongoing management efforts aim to mitigate the effects of FeLV on panthers, but with limited information about which strategies may be most effective and efficient.
2. We used a simulation-based approach to determine optimal FeLV management strategies in panthers. We simulated use of proactive FeLV management strategies (i.e., proactive vaccination) and several reactive strategies, including reactive vaccination and test-and-removal. Vaccination strategies accounted for imperfect vaccine-induced immunity, specifically partial immunity in which all vaccinates achieve partial pathogen protection. We compared the effectiveness of these different strategies in mitigating the number of FeLV mortalities and the duration of outbreaks.
3. Results showed that inadequate proactive vaccination can paradoxically increase the number of disease-induced mortalities in FeLV outbreaks. These effects were most likely due to imperfect vaccine immunity causing vaccinates to serve as a semi-susceptible population, thereby allowing outbreaks to persist in circumstances otherwise conducive to fadeout. Combinations of proactive vaccination with reactive test-and-removal or vaccination, however, had a synergistic effect in reducing impacts of FeLV outbreaks, highlighting the importance of using mixed strategies in pathogen management.
4. *Synthesis and applications:* Management-informed disease simulations are an important tool for identifying unexpected negative consequences and synergies among pathogen management strategies. In particular, we find that imperfect vaccine-induced immunity necessitates further consideration to avoid unintentionally worsening epidemics in some conditions. However, mixing proactive and reactive interventions can improve pathogen control while mitigating uncertainties associated with imperfect interventions.

## Introduction

Outbreaks of infectious diseases can have significant impacts on the population health of free-ranging wildlife and are of heightened importance in species of conservation concern (Breed et al., 2009). A range of pathogen management interventions are available for wildlife, though these options are typically more limited and may be associated with a greater range of uncertainties than in domestic animal or human systems (Miguel et al., 2020). In general, pathogen management strategies can be preventive (hereafter, *proactive*) or reactive. Among proactive and reactive strategies, vaccination is a cornerstone in veterinary medicine (Breed et al., 2009) and functions by reducing the availability of susceptible individuals. Vaccination can be applied randomly, or targeted to: high risk individuals or populations (e.g., Beyer et al., 2012); likely superspreaders (e.g., in chimpanzees: Rushmore et al., 2014); seasonal dynamics (e.g., Baker et al., 2019); or spatial risk, as with vaccine barriers to prevent spread of a pathogen to new areas or populations (e.g., Sanchez & Hudgens, 2020; Slate et al., 2005). However, vaccination comes with myriad uncertainties, including vaccine efficacy, duration of immunity, or even what hosts should be targeted for management of multi-host pathogens (Barnett & Civitello, 2020).

Among reactive pathogen management strategies, test-and-removal (or test-and-cull) is commonly used in domestic and aquaculture species to functionally reduce the infectious period of infected individuals (Miguel et al., 2020). However, this approach is rarely used in wildlife due to the generally low availability of field diagnostic tests and difficulty in recapturing individuals after a positive diagnosis (Miguel et al., 2020). Rather, pathogen control in wildlife often relies on non-selective culling in an attempt to reduce density and—in theory—transmission (Lloyd-Smith et al., 2005). However, non-selective culling can have myriad negative consequences for disease management including removal of immune individuals (Miguel et al., 2020; Potapov et al., 2012) or culling-induced perturbation (e.g., Woodroffe et al., 2006).

While vaccination and culling or removal approaches are just a subset of pathogen management techniques in wildlife, their associated uncertainties are a common feature and hamper efforts to effectively control pathogen transmission. Consequently, mathematical models of infectious disease transmission are a critical tool for optimizing disease management protocols, and have been used in a variety of free-ranging wildlife species of conservation concern, including Ethiopian wolves (Haydon et al., 2006), chimpanzees (Rushmore et al., 2014), and Amur tigers (Gilbert et al., 2020). A key benefit of such models is that they allow the ethical testing of a range of different protocols in light of common management uncertainties such as vaccine efficacy, duration of immunity, and the optimal timing of interventions (Breed et al., 2009). Further, models can serve the important function of balancing the realities of fieldwork with ideal disease control protocols to provide practical, effective guidance for wildlife managers (e.g., Robinson et al., 2018).

Here, we demonstrate the utility of such models for determining optimal disease prevention and control strategies in light of critical management uncertainties for an iconic endangered carnivore, the Florida panther (*Puma concolor coryi*). The feline retrovirus, feline leukemia virus (FeLV), has been the source of significant outbreaks in two endangered felids: Iberian lynx (*Lynx pardinus*) and Florida panthers (*Puma concolor coryi*). In the case of panthers, FeLV caused a deadly outbreak in 2002-2004 (Cunningham et al., 2008), spilling over from domestic cats with subsequent direct transmission among panthers (Brown et al., 2008). In addition, recent evidence demonstrates ongoing FeLV spillover to and transmission among panthers (Chiu et al., 2019), necessitating continued management to prevent future epidemics of this deadly pathogen. FeLV inoculation with a domestic cat vaccine that requires two initial vaccines (boosting) has been used previously in panthers but with unknown efficacy (Cunningham et al., 2008).

Thanks to ready availability of a field diagnostic test, FeLV management in domestic cats and the endangered Iberian lynx has included test-and-removal or isolation of infected individuals (Little et al., 2020; López et al., 2009). Given the use of this intervention in other species and the experienced network of panther veterinarians and rehabilitators in Florida, test-and-isolation (i.e., temporary removal to rehabilitation facilities) or removal is part of future FeLV mitigation plans in panthers. However, uncertainties remain regarding sufficiency of test-and-isolation or removal (hereafter, test-and-removal) to mitigate pathogen spread.

Given the ongoing risks of FeLV to panther conservation and the uncertainties regarding application of proactive and reactive FeLV management strategies, the objectives of this study were to test singly, and in combination, the effectiveness of: (1) proactive vaccination, (2) reactive vaccination, and (3) reactive test-and-removal for reducing the population level impacts of FeLV in Florida panthers. In addition, we evaluated the effectiveness of temporary panther spatial restrictions via closure of wildlife highway underpasses, which we describe and discuss in the supplementary materials. Together, these different strategies and their respective uncertainties represent common tools and considerations for managing pathogens across animal systems, with our results providing extendable insight for veterinarians, researchers, and wildlife managers.

## Methods

### Simulation pipeline

To examine the effect of different disease management regimes on FeLV control, we used spatially explicit network simulations adapted from our research determining drivers of retrovirus transmission in panthers (Gilbertson et al., 2021). This approach involves two steps: (1) simulation of a contact network among panthers, and (2) simulation of FeLV transmission through this network. In brief, we simulated panther populations of 150 individuals (McClintock et al., 2015) and used our previously described exponential random graph model for retrovirus transmission in panthers (Gilbertson et al., 2021) to simulate contact networks constrained by network density among simulated populations (see supplementary methods for further details and Table S1 for parameters).

Briefly, we used a stochastic chain binomial transmission model with a susceptible-infectious-recovered compartmental framework, where individuals could be progressively, regressively, or abortively infected (Cunningham et al., 2008; model developed in Gilbertson et al., 2021). Progressive infections always resulted in death, while regressive infections eventually resolved infection (i.e., recovered with immunity), and abortive infections were always considered immune. Importantly, based on our previous work, we allowed both progressives and regressives to be infectious, though regressives were less likely to transmit (Table S1). The infectious period is long for FeLV (estimated as a mean of 18 weeks, though based on limited observations (Cunningham et al., 2008; see also Gilbertson et al., 2021 and Table S1). We included only disease-induced mortality, and, in order to preserve key network structure, allowed territories vacated by deaths to be reoccupied by new susceptible individuals (hereafter, *respawning*). Outbreaks were initiated by a single, randomly selected non-isolate individual in the population (i.e., one that is connected in the network; see also supplement discussion), and proceeded in weekly time steps for up to five years.

Our primary objective was to examine the effect of different FeLV management regimes on epidemic outcomes, so in our primary simulations, we held network generation and transmission parameters constant at previously supported values (see supplementary methods and Table S1 for further details). We evaluated the consistency of our results to the choice of parameter values with a sensitivity analysis (see below). Hereafter, a *parameter set* represents the unique set of network and transmission parameters for any given set of simulations. A *full simulation* includes simulation of a single contact network and FeLV transmission through that network (with or without management interventions). For each parameter set, we performed 100 full simulations.

For *baseline* and *management scenarios,* we recorded key epidemic outcomes in the absence and presence of interventions, respectively (Figure 1). These key outcomes were (1) the number of mortalities, (2) the duration of an epidemic, and (3) the proportion of epidemics that “failed” per 100 successful epidemics. A failed epidemic (i.e. stochastic fadeout; Craft et al., 2013) was one in which fewer than 5 individuals acquired progressive or regressive infections. As at least 5 individuals acquired progressive infections during the 2002-04 outbreak (Cunningham et al., 2008), this fadeout cutoff allowed for realism in simulated outbreak sizes, while also reducing noise among “successful” epidemics resulting from especially small outbreaks. The outcomes of mortalities and epidemic durations were summarized as median values per parameter set, as results were often skewed; all outcomes were compared between baseline and management scenarios, including quantification with Mann-Whitney *U* tests (supplementary results). All simulations were performed in R version 3.6.3 (R Core Team, 2018).

**Figure 1:**
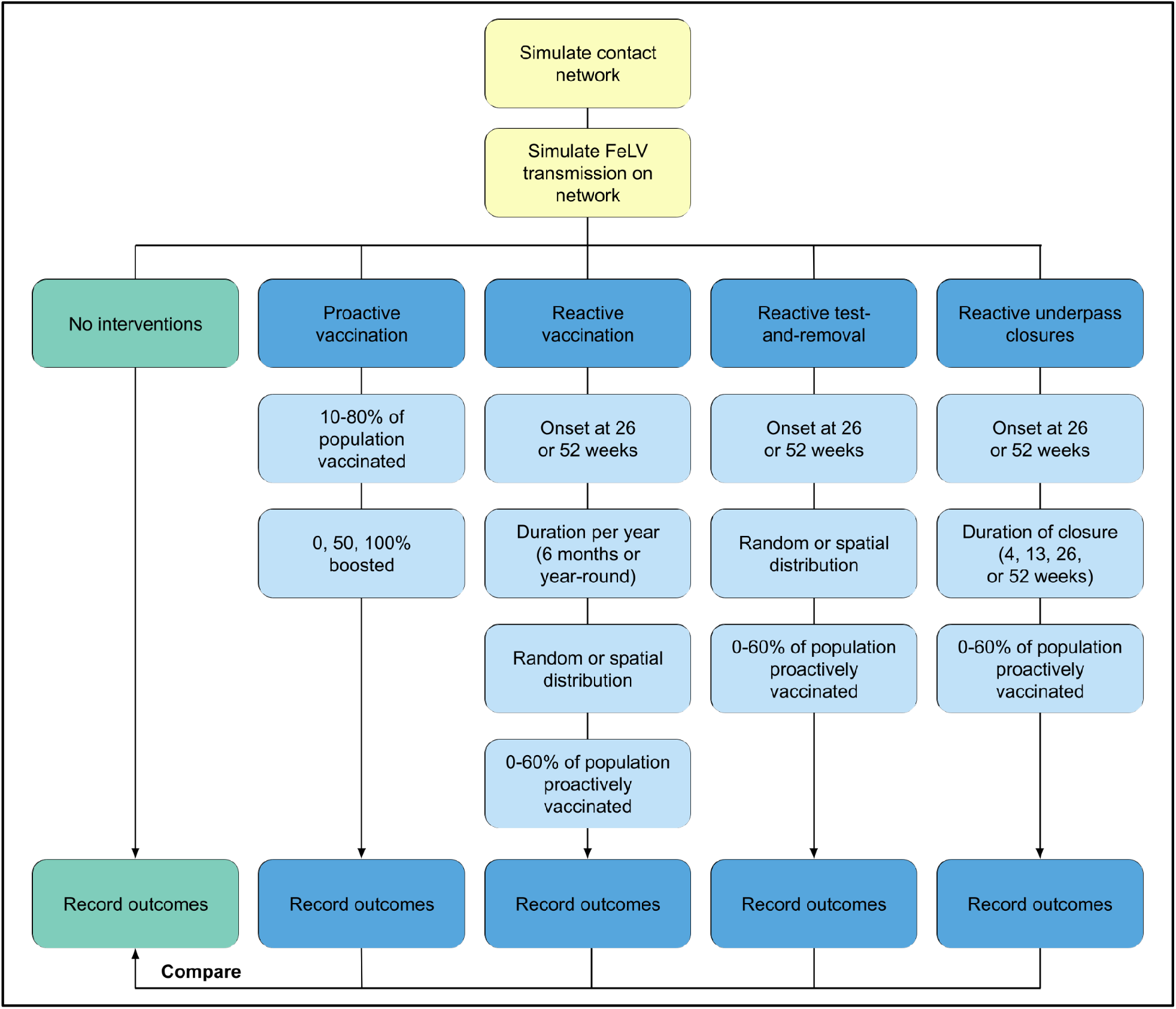
Steps of the simulation process across baseline and management scenarios. The basic network and FeLV transmission simulation steps are shown in yellow, the baseline (no intervention) scenario in green, and the different overarching management scenarios in blue. Within each management scenario, we investigated several variations for the given approach (shown in light blue) in a factorial design. Note that methods and results for reactive underpass closures are described in the supplementary materials.

### Management scenarios

We examined the effect of different levels of population proactive vaccination (proportion of the population vaccinated prior to an outbreak), and different ratios of single versus boosted vaccination. We simulated from 10-80% (in 10% increments) of the population having some degree of vaccine-induced immunity to FeLV prior to the onset of an outbreak. These proactive vaccinations were distributed randomly in the population. Among the vaccinated individuals, 0, 50, or 100% received a second boosting inoculation. Actual vaccine efficacies are unknown for panthers, but based on efficacy studies in domestic cats (reported as preventable fractions; Sparkes, 1997; Torres et al., 2005), we conservatively assumed that boosted vaccination would prevent 80% of infections and single vaccination would prevent 40% of infections. We modeled this efficacy as a binomial probability of protection, wherein increased exposures to infectious individuals increase the likelihood of vaccine failure. Sometimes called “perfectly leaky” immunity, we follow Barnett and Civitello (2020) in referring to this understudied type of imperfect vaccine protection as *partial immunity,* as compared to *binary immunity* (or “all-or-nothing” immunity, wherein some vaccinated individuals are 100% protected and others receive 0% protection; Barnett & Civitello, 2020; Gomes et al., 2014).

In contrast, for reactive vaccination management scenarios, panthers were selected for vaccination *during* outbreaks at a rate of one panther per week. For modeling purposes, we assumed that managers would not know the disease status of an individual selected for vaccination and that vaccination would be ineffective in infectious individuals. In the case of previously vaccinated individuals, re-vaccination changed the vaccine efficacy of singly vaccinated individuals (efficacy of 40%) to the efficacy for boosted individuals (efficacy of 80%). We varied the timing of the onset of reactive vaccination after the initiation of an FeLV outbreak to reflect the difficulty of epidemic detection in this elusive carnivore. We therefore began reactive vaccination at an optimistic, but difficult-to-attain time point of 26 weeks, and a more realistic time point of 52 weeks. In addition, we varied the distribution of reactive vaccination. While proactive vaccines were always distributed randomly, reactive vaccination was either randomly distributed or spatially distributed in an attempted vaccine barrier based on proximity to the I-75 freeway, which runs east-west through primary panther habitat (see supplement for further details). Because vaccination is resource and time intensive, we evaluated the effect of reactive vaccination for 6 months per year versus year-round.

Test-and-removal management scenarios were built around a protocol in which panthers infectious at capture were removed from the population through humane euthanasia or temporary removal until recovery. For simplicity, based on expected fatality of progressive cases, we assumed that all progressively infected individuals would be humanely euthanized at capture, while infectious regressive individuals were temporarily removed from the population until their recovery and re-released into any open territory. A maximum of five individuals were allowed to be temporarily removed in this way at one time based on expected capacity for housing and care.

We expected that managers would be able to capture and test one panther per week at most, and that captures occurred during a 17 week (about 4 month) capture season, in accordance with current panther capture protocols. We assumed that managers would not know the disease status of a target individual until the capture occurred, so captures were not targeted by infection state, and removed individuals were able to transmit prior to removal. We varied the onset of test-and-removal, such that the intervention began 26 or 52 weeks after the initiation of an epidemic. Captures were random or spatially targeted. If spatially targeted, captures (and consequent removals) only occurred on the same side of the I-75 freeway as the initial FeLV infection.

In addition, we simulated temporary spatial segregation of panthers via closure of wildlife highway underpasses along the I-75 freeway, methods and results for which are reported in the supplementary materials. To examine the effects of proactive and reactive management interventions conducted in concert, we included proactive vaccination in all reactive management scenarios: specifically, with 0-60% of the population proactively vaccinated (in 20% increments) prior to initiation of an outbreak. Because it is highly unlikely that 100% of the proactively vaccinated population would have received boosted vaccination, in reactive management scenarios we used a more conservative—yet still challenging to attain—ratio of 50% of the proactively vaccinated population being boosted (i.e. efficacy of 80%). We conducted all management scenarios in a full factorial design resulting in 24, 32, and 16 parameter sets for proactive vaccination, reactive vaccination, and reactive test-and-removal scenarios, respectively (100 full simulations per set for a total of 7,200 simulations).

### Sensitivity analysis

We used a latin hypercube sampling (LHS) approach to generate 50 sensitivity analysis parameter sets across our 8 network and transmission parameters (Table S1) using the *lhs* package in R (Carnell, 2012). We repeated our baseline scenario simulations across these 50 parameter sets and completed 50 simulations per parameter set for all sensitivity analyses.

Due to the high computational effort required to perform sensitivity simulations across all management scenarios, we focused only on the proactive vaccination scenarios. Specifically, we evaluated a subset of 12 LHS parameter sets across a subset of the proactive vaccination conditions in a factorial design (see supplementary methods for further details), resulting in 108 parameter sets with 50 full simulations per parameter set (5,400 full simulations). This approach allowed us to examine sensitivity of our proactive vaccination results across different outbreak sizes and network and transmission parameters, while mitigating computational effort associated with exploring such a wide range of parameters and scenario variations. Sensitivity analysis simulation results were evaluated using scatterplots and Partial Rank Correlation Coefficients (PRCC; Marino et al., 2008; Wu et al., 2013); proactive vaccination scenarios were evaluated for alignment with our qualitative result that low proactive vaccination can increase disease-induced mortalities.

## Results

Progressive infections are expected to result in death in panthers, and are therefore of key concern for management efforts. For simplicity, hereafter, we refer to the number of progressive infections in simulations as the number of mortalities. In the baseline, no-intervention scenario, the median number of mortalities was 34 (range: 1-54); median duration of epidemics was 119.5 weeks, and 34 epidemics failed (fewer than 5 progressive or regressive infections) per 100 successful epidemics.

### Proactive vaccination alone

Proactive vaccination paradoxically increased the number of mortalities across a range of conditions, especially without vaccine boosting (Figures 2, S2; Table S2). Even with 50% of vaccinated individuals receiving a booster, proactive vaccination only reduced mortalities from the baseline scenario at high levels of population vaccination (i.e., 60-80%). With 100% boosting, proactive vaccination increased mortalities at low levels of population vaccination (10-20%), had marginal effects at 30-40% population vaccination, and was strongly effective at about 50% population vaccination levels and higher (at 80% population vaccination, median 17.5 mortalities versus 34 with no interventions). Proactive vaccination consistently lengthened the duration of epidemics, relative to the baseline scenario (up to a median duration of 143.5 weeks; Figures S3-5, Table S3). Of those individuals which became infected in proactive vaccination scenarios, susceptibles were typically infected earliest, followed by vaccinated individuals, and susceptible respawned individuals (Figures 2, S6). When all vaccinated individuals received a booster, vaccination reduced the probability of a successful outbreak even at 40% population vaccination (52 failures versus 34 failures per 100 successful epidemics; Figure S17). In contrast, when no vaccinated individuals received a booster, proactive vaccination was largely only effective at reducing the probability of epidemics at very high levels of population vaccination.

**Figure 2:**
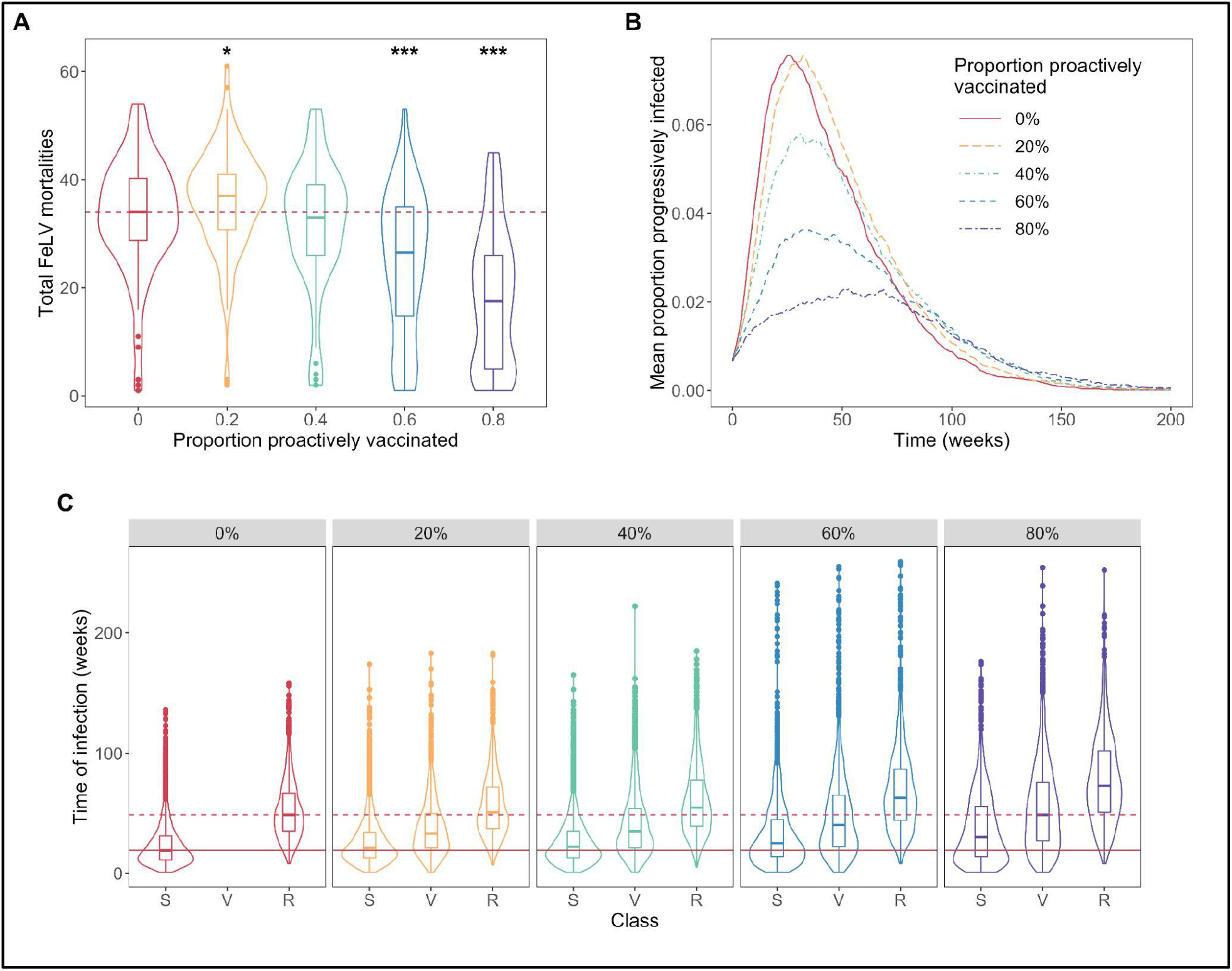
Results of proactive vaccination management scenarios. (A) shows FeLV mortalities with increasing levels of proactive vaccination. The red horizontal line is the median number of mortalities with no interventions. Asterisks indicate statistically significant differences from the no intervention scenario by Mann-Whitney U tests (* <0.05; *** <0.001; Table S2). (B) shows flattening of epidemic curves (mean infections per time step across simulations) with increasing vaccination. (C) shows the time point of infection by class of individual with increasing levels of proactive vaccination, with S = susceptibles, V = vaccinates, and R = respawns. The solid horizontal and dashed lines are the median times of infection for susceptibles and respawns with no interventions, respectively. There is no data for vaccinates in the 0% panel because no individuals were proactively vaccinated. All results are from scenarios in which all vaccinates received a booster.

### Proactive and reactive vaccination

Reactive vaccination alone did not reduce mortalities (Figure 3, Table S4). However, reactive vaccination appeared to work synergistically with proactive vaccination, particularly at moderate to high levels of proactive vaccination (i.e., at least 40-60% of the population proactively vaccinated). This largely held true regardless of the timing of intervention onset and the strategy for reactive vaccination distribution (i.e., random versus spatial; Figures S7-9). The mortality-reducing effects of reactive vaccination were, however, slightly reduced if reactive vaccination occurred for only 6 months out of the year (Figures S8-9). A ratio of greater than 1.5 inoculations per vaccinated individual appeared to promote the largest reductions in mortalities (e.g., as few as a median of 25 mortalties with year-round reactive vaccination and 60% proactive vaccination versus 34 mortalities with no interventions; Figures S7). Adding reactive vaccination largely did not affect the durations of simulated epidemics (Figures S10-11, Table S5), and had little impact on the probability of epidemics failing, particularly in comparison to proactive vaccination alone (Figure S17).

**Figure 3:**
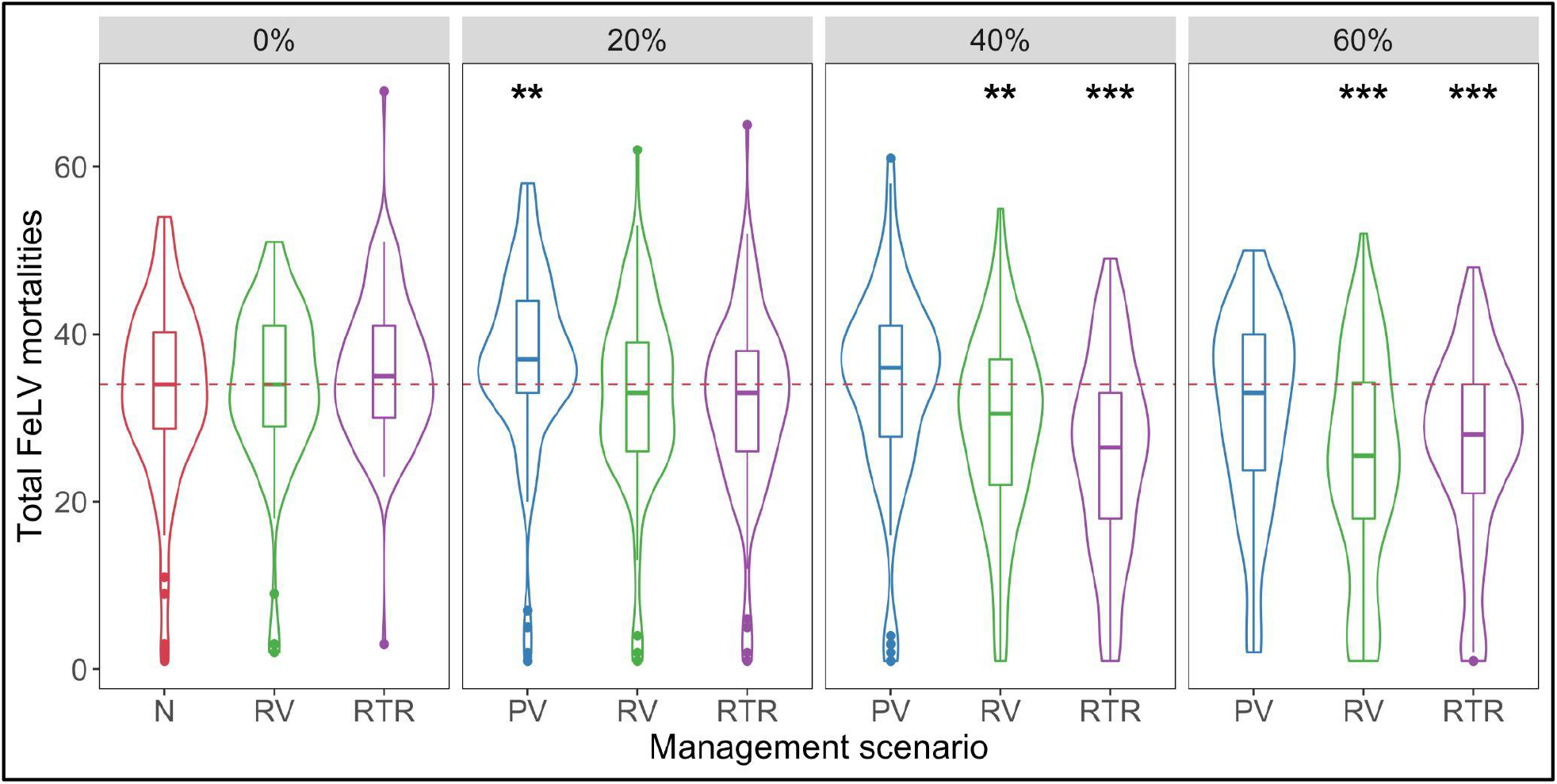
FeLV mortalities with no interventions (N), proactive vaccination alone (PV), reactive vaccination (RV), and reactive test-and-removal (RTR). Because reactive interventions were simulated singly and in combination with proactive vaccination, the individual panels show the level of proactive vaccination (0 – 60%) used with reactive interventions. The horizontal red line gives median mortalties with no interventions (i.e., 0% proactive vaccination). Asterisks indicate statistically significant differences from the no intervention scenario by Mann-Whitney U tests (** <0.01; *** <0.001; Tables S2, S4, S6). All results are for scenarios in which 50% of vaccinates received a booster.

### Proactive vaccination with test-and-removal

Test-and-removal alone did not reduce mortalities (Figure 3, Table S6). Like reactive vaccination, however, test-and-removal appeared to work synergistically with proactive vaccination, especially at moderate to high levels of proactive vaccination (i.e., at least 40-60% of the population proactively vaccinated; as few as a median of 26 mortalities). This largely held true regardless of the timing of the onset of the intervention or the targeting of captures (i.e., random versus spatial). Notably, simulated captures were only conducted for about 4 months per simulation year, in contrast to at least 6 months of reactive vaccination per year. Captures were marginally more likely to successfully identify actively infectious individuals when initiated earlier in an outbreak (Figure S12). The addition of test-and-removal largely did not affect the durations of epidemics (Figure S13, Table S7). When coupled with proactive vaccination, test-and-removal had a modest effect in reducing the probability of a successful epidemic (e.g., maximum of 50 failed epidemics per 100 successful versus 34 failed epidemics per 100 successes with no interventions; Figure S17).

### Sensitivity analyses

Simulated epidemic sizes were variable across the full 50 sensitivity analysis parameter sets in the absence of FeLV interventions (range: median 7.5 mortalities–median 47.5 mortalities; Figure S18). Based on PRCC, the parameters for network density, transmission potential from regressives, weekly contact rates, and baseline transmission potential were positively associated with median mortalities in the absence of interventions; the parameter for the infection-induced mortality rate was negatively associated with median mortalities (see supplementary results, Figure S19).

When focusing on a subset of parameter sets for proactive vaccination sensitivity analysis, low levels of proactive vaccination (e.g., 20% population proactive vaccination) were sometimes effective in reducing the number of mortalities, in contrast to our primary results. PRCC results from proactive scenario sensitivity analysis suggested that the parameters for network density, transmission potential from regressives, and weekly contact rates were positively correlated with increased mortalities at low levels of vaccination (Figure S20). However, network density did not have a clearly monotonic relationship with the difference between mortalities with and without proactive vaccination (Figure S21). We therefore performed additional *post-hoc* sensitivity analyses to further interrogate the relationship between network density and our qualitative outcome of increased mortalities at low levels of proactive vaccination (see supplementary results for detailed discussion of *post-hoc* analysis). These additional analyses found some evidence that low levels of proactive vaccination were least effective at reducing mortalities at intermediate values of parameters governing network connectivity (network density and proportion adults), especially when coupled with increased transmission potential (e.g. higher infectiousness of regressive individuals and/or increased weekly contact rates; Figures S23-24).

## Discussion

In this study we found unexpected consequences and impacts of several epidemic management strategies for disease control in small populations of conservation concern. Our simulation results demonstrate the power of partnering modeling approaches and population management questions to test and optimize disease control strategies in free-ranging wildlife (Joseph et al., 2013). Furthermore, the principles of transmission and pathogen control underlying our findings provide insights for pathogen management in other host-pathogen systems, including humans and livestock.

### Proactive vaccination alone may exacerbate epidemic outcomes under some conditions

Our simulation results showed a paradoxical increase in FeLV mortalities with low levels of proactive vaccination. This counterintuitive finding is likely due, at least in part, to partial vaccine immunity, a type of vaccine imperfection often overlooked in studies of wildlife disease (Barnett & Civitello, 2020). Under partial vaccine immunity, our vaccinated individuals acted as a semi-protected susceptible pool, contracting infection later in the course of an epidemic (Figure 2), thereby acting as a source of new infections later in an outbreak, and ultimately increasing the total number of mortalities. Our sensitivity analysis suggests that increased mortality with inadequate proactive vaccination is a variable result, with populations with intermediate levels of network connectivity and/or high transmission potential being most vulnerable to these paradoxical effects. Transmission in spatially structured populations provides some insight (McCallum, 2008), suggesting that with high or low connectivity, epidemics can fade out quickly; in contrast, with intermediate connectivity, the vaccine failures in our simulations appear to provide a steady supply of new susceptibles to maintain epidemics. Further, in cases of high transmission potential, our results suggest that partial vaccine immunity can shift a rapid fadeout epidemic to a sustained epidemic scenario, as in Rees et al. (2013). While our sensitivity analysis was limited by computational demands and use of some discrete parameters (which may affect PRCC inference; Marino et al., 2008), our findings are consistent with this broader body of literature.

FeLV vaccine efficacy may operate differently in reality from our simulation structure: for example, some vaccinated individuals may have zero vaccine-induced immunity, while others have 100% protection (binary immunity; Barnett & Civitello, 2020). Alternatively, vaccination may not protect from infection but could reduce viral shedding or increase survival of infected individuals (Barnett & Civitello, 2020). In the case of binary immunity, vaccine efficacy would be unlikely to prolong and worsen epidemics as we saw here. In contrast, increased survival of infected individuals without changes to shedding potential could extend or worsen outbreaks, and even favor the evolution of virulence (Barnett & Civitello, 2020). Future research should assess vaccinated panthers’ immune response to FeLV infection *in vitro* or through time dependence of infection in vaccine field trials (Gomes et al., 2014) to refine how imperfect immunity may further alter vaccination guidelines.

Based on our results, we argue that wildlife managers should continue proactively vaccinating animals, as proactive vaccination still frequently reduced mortalities and increased the probability of epidemic failure. In addition, in instances where vaccine boosters provide major gains in vaccine efficacy, our results suggest managers should prioritize boosting vaccinated individuals to develop a core population of high-immunity individuals, rather than a broadly distributed low-immunity population. These recommendations should be most effective at increasing the probability of epidemic failure, and align with other wildlife studies which have emphasized vaccination of core populations (Vial et al., 2006) or risk-based sub-populations (Beyer et al., 2012). Given ongoing uncertainties in vaccine efficacy and the duration of immunity, in the event of an outbreak, we recommend updating model parameterizations to reflect on-the-ground realities in an adaptive management framework in order to provide the most useful predictions and guidance for managers. For example, observed “breakthrough” infection rates (infections in vaccinated individuals) would be a key observation for updating model predictions and adapting management responses.

### Reactive and proactive strategies can work synergistically to reduce epidemic impacts

Our simulations showed that both reactive vaccination and test-and-removal strategies reduced FeLV mortalities in panthers when used in combination with moderate levels of proactive vaccination. Test-and-removal had more consistent effects with arguably less effort than reactive vaccination (4 months of captures compared to year-round reactive vaccination). We therefore suggest that test-and-removal be prioritized over reactive vaccination, especially if identification of actively infectious individuals can be improved.

In our simulations, captures that were most aligned with the initial wave of infectious individuals (i.e., with earlier onset) were more likely to identify actively infectious individuals for removal. This finding highlights the importance of targeting captures to individuals likely to be actively infectious. However, determining infection status in cryptic wildlife is difficult, and consequently supports the increased use of remote tracking technologies that may be able to (1) identify behavior changes associated with sickness, and (2) detect the onset of an epidemic more quickly. This conclusion is consistent with similar findings in Channel Island foxes, where increasing the number and frequency of tracking of sentinel individuals was important for early identification of epidemics (Sanchez & Hudgens, 2020).

A key component of the success of test-and-removal here is the selective removal we simulated, which avoids removing immune individuals that contribute to overall herd immunity (Miguel et al., 2020; Potapov et al., 2012). However, we have simplified the field testing process in our simulations. The common field FeLV diagnostic test identifies antigenemia, which is key for identifying actively infectious individuals. However, the duration of antigenemia—and even degree of infectiousness—in regressively infected individuals is unclear in panthers. We may therefore overestimate the effect of removing regressive individuals, but given their reduced infectiousness in our simulations, we still expect test-and-removal to be a key strategy for mitigating FeLV impacts in panthers.

Notably, reactive vaccination, in concert with proactive vaccination, is also a viable alternative strategy to test-and-removal. Our simulations showed reactive vaccination to have the strongest effects for reducing FeLV mortalities when at least 50% of the population was vaccinated, and with a ratio of about 1.5 vaccines per vaccinated individual (i.e., 50% of vaccinated individuals received a booster). We therefore suggest that managers should prioritize boosting at least half of vaccinated individuals in a reactive vaccination response scenario. We also note that without ongoing vaccination (i.e., reactive vaccination), births can provide an additional pool of susceptibles that may maintain epidemics (births were functionally represented by respawns in our simulations). Reactive vaccination can thus be critical for preventing a shift to endemicity among invading pathogens.

We did not see a strong effect of attempting a vaccine barrier, in contrast to Sanchez et al. (2020), where a simulated vaccine barrier could effectively halt spread of a pathogen in Channel Island foxes. This difference is likely due to the differences in home range size and movement capacity between the two species, with foxes ranging less widely than panthers. In reality—and as part of an adaptive management response—if an empirical pathogen outbreak exhibited a stronger spatial signal than was featured in our simulations, spatially targeted reactive vaccination would still be a worthwhile intervention strategy.

Importantly, both reactive vaccination and test-and-removal strategies mitigated the negative effects of low levels of population protection seen with inadequate proactive vaccination. Particularly if levels of population protection from proactive vaccination are unknown, it is vital for managers to incorporate these reactive strategies for pathogen management.

### Limitations and future directions

In this study, we considered the effects of partial vaccine immunity, but we made the simplifying assumption of no waning vaccine or infection-induced immunity over time. However, our findings with regard to imperfect efficacy are representative of the likely consequences of waning immunity in that the loss of immunity supplies new susceptibles to the population. Should immunity not outlast the course of an FeLV outbreak, this process could prolong outbreaks and result in increased mortalities. We therefore recommend that future research examine the effects of waning vaccine immunity, particularly considering the value of revaccinating individuals that may be experiencing loss of vaccine protection. Furthermore, an adaptive management framework should monitor for infection in individuals previously considered immune, which could represent vaccine failure or fading immunity. Model predictions could then be updated with these observations and revaccination rates adjusted accordingly.

Our simulation results found that relatively high levels of vaccination were required to reduce impacts of FeLV in panthers, in contrast to studies in other endangered species (Gilbert et al., 2020; Haydon et al., 2006). However, here we investigated a pathogen with a long duration of infectiousness and lacked the advantages of distinct corridors between panther subpopulations for reducing required levels of vaccination. It is therefore unsurprising that panthers would require higher levels of FeLV population vaccination than was found, for example, for rabies vaccination in Ethiopian wolves (Haydon et al., 2006) or canine distemper virus vaccination in Amur tigers (Gilbert et al., 2020). However, we also did not assume the presence of preexisting population immunity prior to proactive vaccination, and some degree of population immunity likely already exists in panthers, given ongoing exposures (Chiu et al., 2019). This would reduce necessary vaccination levels in panthers, as would higher vaccine efficacy than we conservatively assumed here (Vial et al., 2006). We further point out that our results should not be treated as scenario-specific predictions; rather, our findings should be considered in terms of the consistency of trends across parameter variations in both our main simulations and sensitivity analyses.

### Conclusions

Our simulation results highlight the benefits of using a mixture of proactive and reactive interventions in pathogen management, particularly in the context of uncertain vaccine efficacy or population protection. Further, vaccine studies should investigate the mechanisms underlying imperfect immunity, with modeling studies a useful tool for estimating how imperfect immunity— whether binary or partial—affect consequent management recommendations. This research highlights the value of linking modeling and management priorities to identify unexpected consequences of interventions and determine optimal pathogen management strategies in free-ranging wildlife.

## Authors’ Contributions

All authors conceived the ideas and designed methodology; MLJG performed simulations, analyzed data, and led the writing of the manuscript. All authors contributed critically to the drafts and gave final approval for publication.

## Acknowledgements

This research was supported by the National Science Foundation (DEB-1413925, 1654609, and 2030509). MLJG was supported by the Office of the Director, National Institutes of Health (NIH T32OD010993), the University of Minnesota Informatics Institute MnDRIVE program, and the Van Sloun Foundation. The content is solely the responsibility of the authors and does not necessarily represent the official views of the National Institutes of Health. Florida panther data collected by the Florida Fish and Wildlife Conservation Commission is fully supported by donations to the Florida Panther Research and Management Trust Fund via the registration of “Protect the Panther’’ license plates. We acknowledge the efforts of National Park Service staff in the collection of Florida panther data utilized in this study.

## Data Availability

Full R code for simulations is available on GitHub (https://github.com/miones029/FeLV_Management_Simulations) and upon acceptance will be archived at Zenodo.

## Works Cited

Baker, L., Matthiopoulos, J., Müller, T., Freuling, C., & Hampson, K. (2019). Optimizing spatial and seasonal deployment of vaccination campaigns to eliminate wildlife rabies. Philosophical Transactions of the Royal Society of London. Series B, Biological Sciences, 374(1776), 20180280.

Barnett, K. M., & Civitello, D. J. (2020). Ecological and Evolutionary Challenges for Wildlife Vaccination. Trends in Parasitology, 36(12), 970–978.

Beyer, H. L., Hampson, K., Lembo, T., Cleaveland, S., Kaare, M., & Haydon, D. T. (2012). The implications of metapopulation dynamics on the design of vaccination campaigns. Vaccine, 30(6), 1014–1022.

Breed, A. C., Plowright, R. K., Hayman, D. T. S., Knobel, D. L., Molenaar, F. M., Gardner-Roberts, D., Cleaveland, S., Haydon, D. T., Kock, R. A., Cunningham, A. A., Sainsbury, A. W., & Delahay, R. J. (2009). Disease Management in Endangered Mammals. In R. J. Delahay, G. C. Smith, & M. R. Hutchings (Eds.), Management of Disease in Wild Mammals (pp. 215–239). Springer Japan.

Brown, M. A., Cunningham, M. W., Roca, A. L., Troyer, J. L., Johnson, W. E., & O’Brien, S. J. (2008). Genetic characterization of feline leukemia virus from Florida panthers. Emerging Infectious Diseases, 14(2), 252–259.

Carnell, R. (2012). lhs: Latin hypercube samples. R Package Version 0.10, URL http://CRAN.R-Project.org/package=Lhs.

Chiu, E. S., Kraberger, S., Cunningham, M., Cusack, L., Roelke, M., & VandeWoude, S. (2019). Multiple Introductions of Domestic Cat Feline Leukemia Virus in Endangered Florida Panthers. Emerging Infectious Diseases, 25(1), 92–101.

Craft, M. E., Beyer, H. L., & Haydon, D. T. (2013). Estimating the probability of a major outbreak from the timing of early cases: an indeterminate problem? PloS One, 8(3), e57878.

Cunningham, M. W., Brown, M. A., Shindle, D. B., Terrell, S. P., Hayes, K. A., Ferree, B. C., McBride, R. T., Blankenship, E. L., Jansen, D., Citino, S. B., Roelke, M. E., Kiltie, R. A., Troyer, J. L., & O’Brien, S. J. (2008). Epizootiology and management of feline leukemia virus in the Florida puma. Journal of Wildlife Diseases, 44(3), 537–552.

Gilbert, M., Sulikhan, N., Uphyrkina, O., Goncharuk, M., Kerley, L., Hernandez Castro, E., Reeve, R., Seimon, T., McAloose, D., Seryodkin, I. V., Naidenko, S. V., Davis, C. A., Wilkie, G. S., Vattipally, S. B., Adamson, W. E., Hinds, C., Thomson, E. C., Willett, B. J., Hosie, M. J. … Cleaveland, S. (2020). Distemper, extinction, and vaccination of the Amur tiger. Proceedings of the National Academy of Sciences of the United States of America, 117(50), 31954–31962. https://doi.org/10.1073/pnas.2000153117

Gilbertson, M. L. J., Fountain-Jones, N. M., Malmberg, J. L., Gagne, R. B., Lee, J. S., Kraberger, S., Kechejian, S., Petch, R., Chiu, E., Onorato, D., Cunningham, M. W., Crooks, K. R., Funk, W. C., Carver, S., VandeWoude, S., VanderWaal, K., & Craft, M. E. (2021). Transmission of one predicts another: Apathogenic proxies for transmission dynamics of a fatal virus. bioRxiv. https://doi.org/10.1101/2021.01.09.426055

Gomes, M. G. M., Lipsitch, M., Wargo, A. R., Kurath, G., Rebelo, C., Medley, G. F., & Coutinho, A. (2014). A missing dimension in measures of vaccination impacts. PLoS Pathogens, 10(3), e1003849.

Haydon, D. T., Randall, D. A., Matthews, L., Knobel, D. L., Tallents, L. A., Gravenor, M. B., Williams, S. D., Pollinger, J. P., Cleaveland, S., Woolhouse, M. E. J., Sillero-Zubiri, C., Marino, J., Macdonald, D. W., & Laurenson, M. K. (2006). Low-coverage vaccination strategies for the conservation of endangered species. Nature, 443(7112), 692–695.

Joseph, M. B., Mihaljevic, J. R., Arellano, A. L., Kueneman, J. G., Preston, D. L., Cross, P. C., & Johnson, P. T. J. (2013). Taming wildlife disease: bridging the gap between science and management. The Journal of Applied Ecology, 50(3), 702–712.

Little, S., Levy, J., Hartmann, K., Hofmann-Lehmann, R., Hosie, M., Olah, G., & Denis, K. S. (2020). 2020 AAFP Feline Retrovirus Testing and Management Guidelines. Journal of Feline Medicine and Surgery, 22(1), 5–30.

Lloyd-Smith, J. O., Cross, P. C., Briggs, C. J., Daugherty, M., Getz, W. M., Latto, J., Sanchez, M. S., Smith, A. B., & Swei, A. (2005). Should we expect population thresholds for wildlife disease? Trends in Ecology & Evolution, 20(9), 511–519.

López, G., López-Parra, M., Fernández, L., Martínez-Granados, C., Martínez, F., Meli, M. L., Gil-Sánchez, J. M., Viqueira, N., Díaz-Portero, M. A., Cadenas, R., Lutz, H., Vargas, A., & Simón, M. A. (2009). Management measures to control a feline leukemia virus outbreak in the endangered Iberian lynx. Animal Conservation, 12(3), 173–182.

Marino, S., Hogue, I. B., Ray, C. J., & Kirschner, D. E. (2008). A methodology for performing global uncertainty and sensitivity analysis in systems biology. Journal of Theoretical Biology, 254(1), 178–196.

McCallum, H. (2008). Landscape structure, disturbance, and disease dynamics. Infectious Disease Ecology: Effects of Ecosystems on Disease and of Disease on Ecosystems, 100–122.

McClintock, B. T., Onorato, D. P., & Martin, J. (2015). Endangered Florida panther population size determined from public reports of motor vehicle collision mortalities. The Journal of Applied Ecology, 52(4), 893–901.

Miguel, E., Grosbois, V., Caron, A., Pople, D., Roche, B., & Donnelly, C. A. (2020). A systemic approach to assess the potential and risks of wildlife culling for infectious disease control. Communications Biology, 3(1), 353.

Potapov, A., Merrill, E., & Lewis, M. A. (2012). Wildlife disease elimination and density dependence. Proceedings. Biological Sciences / The Royal Society, 279(1741), 3139–3145.

R Core Team. (2018). R: A Language and Environment for Statistical Computing. R Foundation for Statistical Computing. https://www.R-project.org/

Rees, E. E., Pond, B. A., Tinline, R. R., & Bélanger, D. (2013). Modelling the effect of landscape heterogeneity on the efficacy of vaccination for wildlife infectious disease control. The Journal of Applied Ecology, 50(4), 881–891.

Robinson, S. J., Barbieri M. M., Murphy S., Baker J. D., Harting A. L., Craft M. E., & Littnan C. L. (2018). Model recommendations meet management reality: implementation and evaluation of a network-informed vaccination effort for endangered Hawaiian monk seals. Proceedings of the Royal Society B: Biological Sciences, 285(1870), 20171899.

Rushmore, J., Caillaud, D., Hall, R. J., Stumpf, R. M., Meyers, L. A., & Altizer, S. (2014). Network-based vaccination improves prospects for disease control in wild chimpanzees. Journal of the Royal Society, Interface / the Royal Society, 11(97), 20140349.

Sanchez, J. N., & Hudgens, B. R. (2020). Vaccination and monitoring strategies for epidemic prevention and detection in the Channel Island fox (*Urocyon littoralis*). PloS One, 15(5), e0232705.

Slate, D., Rupprecht, C. E., Rooney, J. A., Donovan, D., Lein, D. H., & Chipman, R. B. (2005). Status of oral rabies vaccination in wild carnivores in the United States. Virus Research, 111(1), 68–76.

Sparkes, A. H. (1997). Feline leukaemia virus: a review of immunity and vaccination. The Journal of Small Animal Practice, 38(5), 187–194.

Torres, A. N., Mathiason, C. K., & Hoover, E. A. (2005). Re-examination of feline leukemia virus: host relationships using real-time PCR. Virology, 332(1), 272–283.

Vial, F., Cleaveland, S., Rasmussen, G., & Haydon, D. T. (2006). Development of vaccination strategies for the management of rabies in African wild dogs. Biological Conservation, 131(2), 180–192.

Woodroffe, R., Donnelly, C. A., & Cox, D. R. (2006). Effects of culling on badger *Meles meles* spatial organization: implications for the control of bovine tuberculosis. Journal of Applied Ecology. 43(1), 1–10. https://doi.org/10.1111/j.1365-2664.2005.01144.x

Wu, J., Dhingra, R., Gambhir, M., & Remais, J. V. (2013). Sensitivity analysis of infectious disease models: methods, advances and their application. Journal of the Royal Society, Interface / the Royal Society, 10(86), 20121018.

