## Supplementary Materials for "Paradoxes and synergies: optimizing management of a deadly virus in an endangered carnivore"

---

---

### Methods

*Network and transmission simulations*

The following is summarized from Gilbertson et al. (2021) where additional details about network and transmission simulations can be found.

We used a spatially-explicit network simulation approach for our simulations, as this allowed us to incorporate both our previously identified drivers of retrovirus transmission and contact heterogeneity important for shaping epidemic outcomes (Keeling & Eames, 2005; Lloyd-Smith et al., 2005). More specifically, we used our previously defined exponential random graph model (ERGM) for retrovirus transmission pathways in panthers to simulate contact networks. Networks were therefore simulated based on pairwise geographic distances between panther home range centroids, proportion of the population that is adult (versus subadult), and two network structural terms: alternating k-stars and geometrically weighted edgewise shared partner distribution. To generate these networks, we first simulated panther populations with these key characteristics (i.e. age classes and distances), and then used ERGM coefficients from Gilbertson et al. (2021) to generate contact networks representing likely FeLV transmission pathways.

We used empirical telemetry data to simulate pairwise geographic distances among each simulated panther population. Panthers have been monitored using predominantly VHF telemetry for decades, with aerial relocations recorded typically every 3 days. The number of collared individuals has decreased in recent years, but as we were focused on FeLV management in the contemporary population (after the 2002-2004 fatal FeLV outbreak), we focused our telemetry data analysis on the most recent years with consistently high relative telemetry coverage (2010-2012; 56 total panthers monitored, with an average of 34 monitored per year). This telemetry data was then used to simulate home range centroids and subsequent pairwise distances.

To simulate age distributions, we randomly assigned age categories (adult versus subadult) based on a “proportion adult” parameter (Table S1, “Adult_prop”), taken from random forest analysis. In simulating networks, we also constrained network density based on target values again identified in random forest analysis (Table S1, “Net_dens”). Similarly, for simulating FeLV transmission along networks, we used a susceptible-infectious-recovered framework and parameterization adapted random forest results (Table S1, Figure S1).

We held all network and transmission parameters constant for our primary baseline (no-intervention) and management scenarios to evaluate only the effect of interventions (Table S1). For sensitivity analysis, we considered the same reasonable parameter ranges used in Gilbertson et al. (2021; Table S1).

**Table S1: Network and transmission simulation parameters**

| Parameter | Definition | Target Value | Range |
|---|---|---|---|
| Adult_prop | Proportion adults versus subadults | 0.89 | 0.82-0.99 |
| Net_dens | Simulated network density | 0.08 | 0.05-0.15 |
| $\beta$ | Probability of transmission from progressives, given effective contact | 0.27 | 0.17-0.29 |
| $C$ | Constant multiplier for probability of transmission from regressives, given effective contact | 0.1 | 0, 0.1, 0.5 |
| $\omega$ | Weekly probability of contact | 0.3 | 0.1-0.4 |

| | | | |
|---|---|---|---|
| $\mu$ | Weekly probability of death from progressive infection | 1/18 | 1/18, 1/26 |
| $K$ | Constant multiplier for weekly probability of recovery from regressive infection | 1 | 0.5, 1 |
| $\nu$ | Weekly probability of territory repopulation ("respawn rate") | 0.11 | 0.083-0.25 |
| $P$ | Proportion randomly assigned to progressive, regressive | 0.25 | 0.25 |

*Note: Values and ranges adapted from Gilbertson et al. (2021). Target values are those used in no-intervention and management scenarios. The ranges column gives the range of possible values used in the Latin hypercube sampling sensitivity analysis (see below). See Figure S1 for transmission parameters in the context of the compartmental model.*

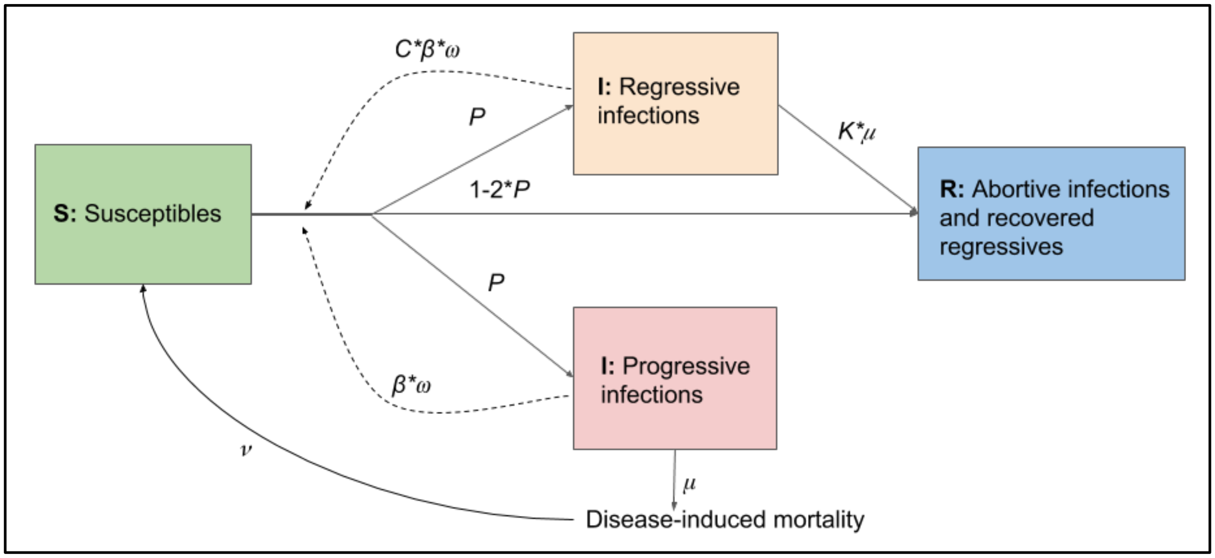

**Figure S1:** Compartmental model diagram of FeLV transmission model (see Table S1 for parameter definitions). Figure adapted from Gilbertson et al. (2021). Note that, for simplicity, the vaccinated class of individuals is not shown.

*Spatial distribution of reactive vaccination*

To simulate spatially-targeted vaccination in a vaccine barrier along the I-75 freeway, we selected individuals for attempted vaccination based on their proximity to the freeway. Proximity was determined by the distance between each individual's simulated home range centroid and latitude 26.16 (representing I-75). Proximities were ranked into three categories (i.e., closest, middle, and furthest thirds of the population), and weighted for sampling. The closest third received a weight of 0.7; the middle third, 0.2; and the furthest third, 0.1. Weighting allowed us to preferentially select individuals with home range centroids close to I-75, but also represented the real-world possibility that panthers range widely and may be identified further from these centroids. Weighting also accounted for the fact that, as individuals close to the freeway were "saturated" with vaccination, other, more distant individuals might become more available candidates for vaccination.

*Reactive underpass closures*

Large-scale isolation or quarantine measures may also be used for disease management in free-ranging wildlife in the form of physical or behavioral barriers between affected and unaffected subsets of a population. For example, fencing has been used to prevent transmission of foot-and-mouth disease between wildlife and cattle in South Africa (Mysterud & Rolandsen, 2019), and winter feeding grounds may help prevent transmission of brucellosis between elk and cattle in the Greater Yellowstone Ecosystem through behavioral separation (Cotterill et al., 2018). For panthers, while physical barriers to prevent spillover from domestic cats are impractical, it may be possible to use temporary spatial barriers to reduce the spatial spread of FeLV among panthers after a spillover event. Specifically, the major I-75 freeway is fenced throughout Florida panther habitat to reduce vehicle strikes, with regular wildlife underpasses the main means for wildlife to traverse this barrier. While never before attempted, it may be possible to physically block these underpasses under emergency conditions to prevent the spread of FeLV from the northern to southern subsets of the panther population, or vice versa.

To determine the potential utility of closing I-75 wildlife underpasses in a FeLV outbreak, we simulated blockage of these underpasses in the same simulation framework described for management scenarios in the main text. We considered a “best case” scenario in which underpass closures were completely effective at preventing transmission across the freeway. Underpasses were closed instantaneously either 26 or 52 weeks after the initiation of an FeLV outbreak, and remained closed for 4, 13, 26, or 52 weeks. We again included variations in proactive vaccination (0-60% in 20% increments), as in other reactive management scenarios. We used a factorial design across the management variations we considered which resulted in 32 parameter sets for underpass closure scenarios (a total of 3,200 full simulations).

*Sensitivity analyses supplementary methods*

To mitigate computational complexity for evaluating sensitivity of our proactive vaccination results and to cover a wide range of parameter space, we focused on the subset of proactive vaccination conditions in which 20, 40, or 60% of the population was proactively vaccinated, with 0, 50, or 100% of vaccinates receiving a booster. Furthermore, we selected a subset of LHS parameter sets for these proactive vaccination sensitivity tests, choosing these based on the results of the full sensitivity analysis under the baseline scenario (see main text). There, we recorded the median number of progressive infections for each of the 50 parameter sets, then randomly selected sets based on quantiles (3 parameter sets from the lower quantile, 6 from the interquartile range, and 3 from the upper quantile). As reported in the main text, this approach allowed us to examine sensitivity of our proactive vaccination results across a wide range of outbreak sizes and network and transmission parameters, while mitigating computational effort associated with exploring such a wide range of parameters and scenario variations. The resulting 12 LHS parameter sets, used in a factorial design across proactive vaccination conditions, resulted in 108 parameter sets with 50 full simulations per set (5,400 full simulations).

To evaluate sensitivity analysis simulation results, we used a combination of scatterplots and Partial Rank Correlation Coefficients (PRCC; Marino et al., 2008; Wu et al., 2013). For baseline, no intervention scenarios, we tested for parameters of significance for the outcome of median mortalities per parameter set. We used averaged (median) mortalities to both minimize the effect of aleatory uncertainty and increase statistical power of PRCC (Marino et al., 2008).

For proactive vaccination sensitivity analyses, we first determined if simulation results were consistent with our qualitative outcome that low vaccination levels can increase mortalities using histograms and median mortalities as in the main text. Finding that this result was not consistent across all parameter sets, we performed additional *post-hoc* analyses and simulations to determine parameters important for increased mortalities at low vaccination levels (see supplementary results below).

**Results**

*Management scenario supplementary results*

For transparency, we present the results for all management scenarios here, including quantification of differences in epidemic outcomes as performed using Mann-Whitney $U$ tests (comparing a given outcome between baseline and management scenarios). Full results for proactive vaccination alone are shown in Figure S2 and Table S2 (mortalities) and Figure S3 and Table S3 (epidemic durations). In addition, we show the relationship between outbreak duration and number of mortalities in Figure S4, demonstrating that longer outbreaks are typically associated with higher numbers of mortalities. Figure S5 shows epidemic curve summaries across a range of proactive vaccination levels, demonstrating how the shape of epidemic curves becomes “flattened” with the use of proactive vaccination. In Figure S6, we show how proactive vaccination interacts with our respawning process to push the time of infection later as proactive vaccination levels are increased. Full results for reactive vaccination mortalities are shown in Figures S7-9 and Table S4; durations in Figure S10-11 and Table S5. Full results for reactive test-and-removal are shown in Figure S12 and Table S6 (mortalities), and Figure S13 and Table S7 (epidemic durations). Full results for reactive underpass closures are reported below (see "Reactive underpass closures").

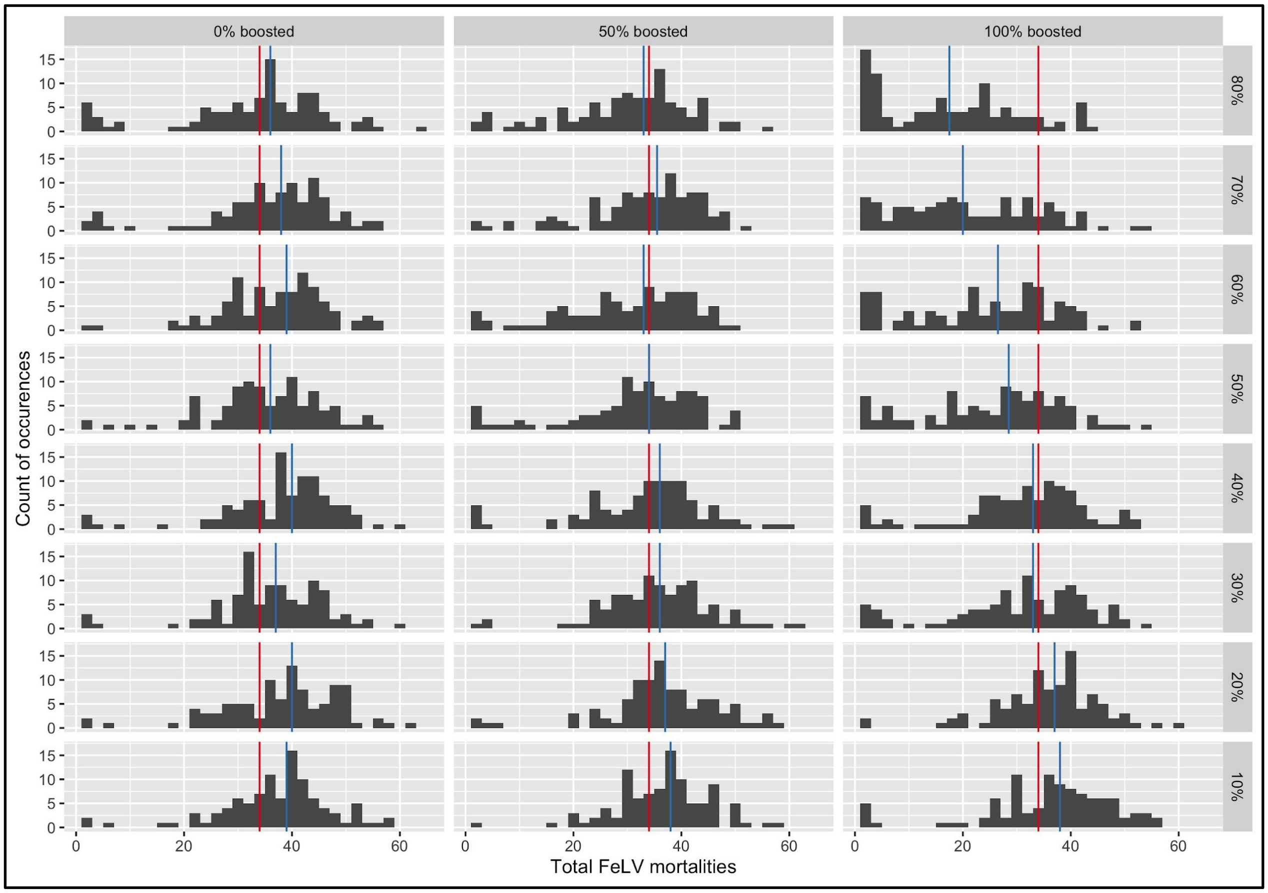

**Figure S2:** Histograms of FeLV mortalities in simulated epidemics with proactive vaccination alone. Red vertical lines indicate the median number of mortalities from the baseline scenario without interventions; blue lines indicate median number of mortalities with proactive vaccination. Panel rows represent the proportion of the population proactively vaccinated; columns represent ratios of vaccine efficacy among vaccinates. Vaccination without a booster was assumed to have 40% efficacy, versus 80% efficacy with a boosting inoculation. Each histogram plot represents the results of 100 simulations.

**Table S2: Mann-Whitney *U* results for mortalities under proactive vaccination**

| Proportion proactive | Proportion boosted | MW test statistic | Estimate | 95% CI (L) | 95% CI (U) | *p-value* |
|---|---|---|---|---|---|---|
| 10% | 0% boosted | 6318 | 4 | 2 | 7 | **<0.01** |
| 20% | 0% boosted | 6482 | 5 | 3 | 8 | **<0.001** |
| 30% | 0% boosted | 5964 | 3 | 1 | 6 | **0.02** |
| 40% | 0% boosted | 6647 | 5 | 3 | 8 | **<0.001** |
| 50% | 0% boosted | 5647 | 2 | -1 | 5 | 0.11 |
| 60% | 0% boosted | 6153 | 4 | 1 | 6 | **<0.01** |
| 70% | 0% boosted | 6051 | 4 | 1 | 6 | **0.01** |
| 80% | 0% boosted | 5325 | 1 | -2 | 4 | 0.43 |
| 10% | 50% boosted | 6057 | 3 | 1 | 6 | **<0.01** |
| 20% | 50% boosted | 6107 | 4 | 1 | 6 | **<0.01** |
| 30% | 50% boosted | 5695 | 2 | 0 | 5 | 0.09 |
| 40% | 50% boosted | 5270 | 1 | -2 | 4 | 0.51 |
| 50% | 50% boosted | 4805 | -1 | -3 | 2 | 0.63 |
| 60% | 50% boosted | 4450 | -2 | -5 | 1 | 0.18 |
| 70% | 50% boosted | 5278 | 1 | -2 | 4 | 0.5 |
| 80% | 50% boosted | 4373 | -2 | -5 | 1 | 0.13 |
| 10% | 100% boosted | 6062 | 4 | 1 | 7 | **<0.01** |
| 20% | 100% boosted | 5878 | 3 | 0 | 5 | **0.03** |
| 30% | 100% boosted | 4621 | -1 | -4 | 2 | 0.35 |
| 40% | 100% boosted | 4589 | -1 | -4 | 1 | 0.32 |
| 50% | 100% boosted | 3351 | -7 | -10 | -3 | **<0.001** |
| 60% | 100% boosted | 3108 | -8 | -12 | -5 | **<0.001** |

| | | | | | | |
|---|---|---|---|---|---|---|
| 70% | 100% boosted | 2501 | -12 | -16 | -8 | **<0.001** |
| 80% | 100% boosted | 1838 | -16 | -20 | -12 | **<0.001** |

*Note: All tests are comparing the given scenario for proactive vaccination (table rows) against the baseline scenario. Statistically significant results are highlighted by bold text for* p-values*. Colors indicate the direction of change: red indicates that proactive vaccination significantly increased mortalities, and blue that proactive vaccination significantly reduced mortalities. The proportion proactive column gives the proportion of population proactively vaccinated; proportion boosted is the proportion of those vaccinated which received a second, boosting vaccination. MW test statistic is the Mann-Whitney* U *test statistic. The Estimate column gives the Mann-Whitney* U *difference estimate, and the 95% confidence interval is given in the 95% CI (L) (lower) and 95% CI (U) (upper) columns.*

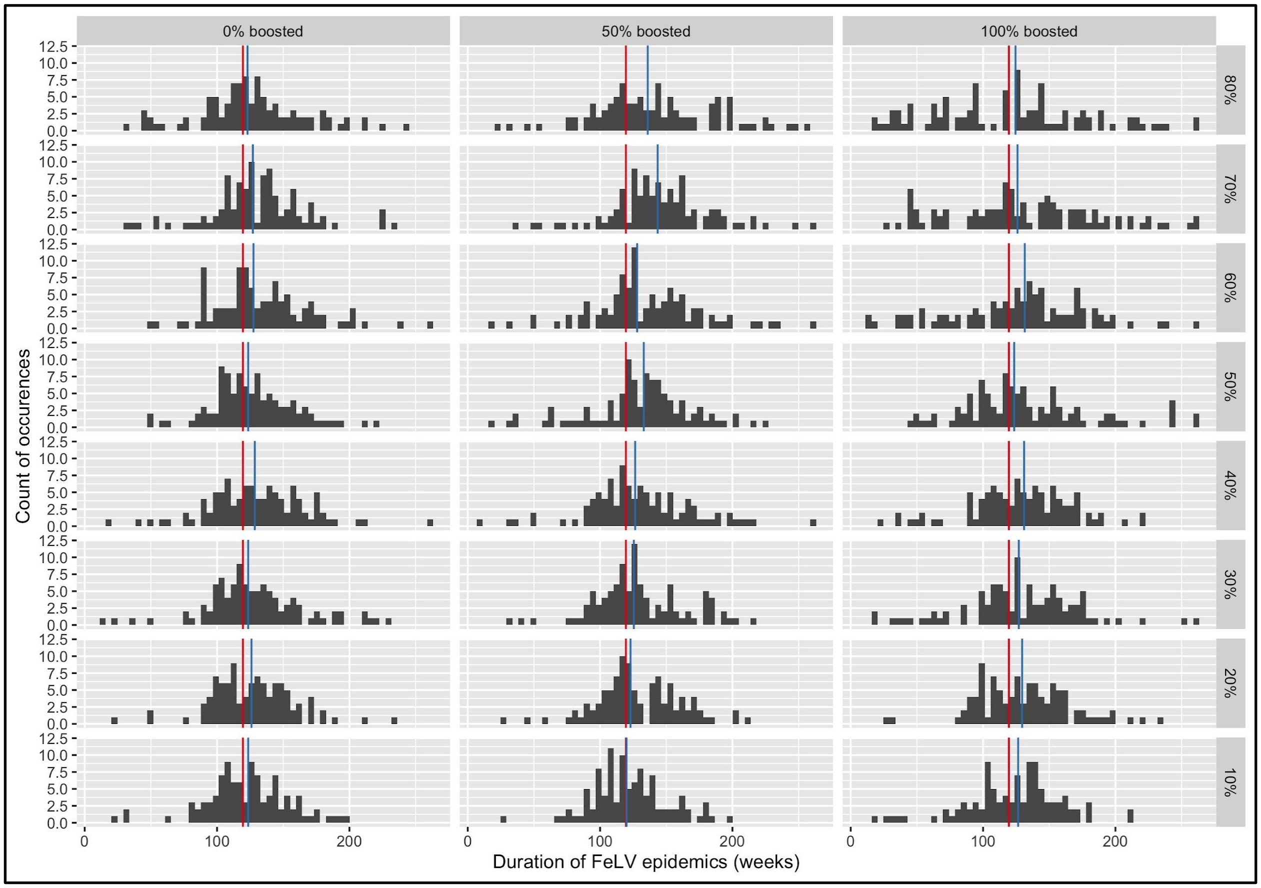

**Figure S3:** Histograms of the duration of simulated FeLV epidemics (in weeks) with proactive vaccination alone. Red vertical lines indicate the median duration from the baseline scenario without interventions; blue lines indicate median duration with proactive vaccination. Panel rows represent the proportion of the population proactively vaccinated; columns represent ratios of vaccine efficacy among vaccinates. Vaccination without a booster was assumed to have 40% efficacy, versus 80% efficacy with a boosting inoculation. Each histogram plot represents the results of 100 simulations.

**Table S3: Mann-Whitney *U* results for epidemic durations under proactive vaccination**

| Proportion proactive | Proportion boosted | MW test statistic | Estimate | 95% CI (L) | 95% CI (U) | *p-value* |
|---|---|---|---|---|---|---|
| 10% | 0% boosted | 5172 | 2 | -6 | 10 | 0.68 |
| 20% | 0% boosted | 5485 | 5 | -3 | 13 | 0.24 |
| 30% | 0% boosted | 5494 | 5 | -3 | 14 | 0.23 |
| 40% | 0% boosted | 5719 | 8 | -1 | 17 | 0.08 |
| 50% | 0% boosted | 5472 | 5 | -4 | 13 | 0.25 |
| 60% | 0% boosted | 5901 | 10 | 1 | 19 | **0.03** |
| 70% | 0% boosted | 5773 | 9 | 0 | 17 | 0.06 |
| 80% | 0% boosted | 5396 | 4 | -5 | 14 | 0.33 |
| 10% | 50% boosted | 5063 | 1 | -8 | 9 | 0.88 |
| 20% | 50% boosted | 5663 | 7 | -1 | 15 | 0.11 |
| 30% | 50% boosted | 5626 | 7 | -2 | 15 | 0.13 |
| 40% | 50% boosted | 5687 | 7 | -2 | 17 | 0.09 |
| 50% | 50% boosted | 5967 | 11 | 2 | 20 | **0.02** |
| 60% | 50% boosted | 6093 | 13 | 3 | 21 | **<0.01** |
| 70% | 50% boosted | 6950 | 22 | 13 | 31 | **<0.001** |
| 80% | 50% boosted | 6278 | 16 | 6 | 27 | **<0.01** |
| 10% | 100% boosted | 5238 | 3 | -6 | 12 | 0.56 |
| 20% | 100% boosted | 5798 | 9 | 0 | 18 | 0.051 |

| | | | | | | |
|---|---|---|---|---|---|---|
| 30% | 100% boosted | 5616 | 7 | -2 | 17 | 0.13 |
| 40% | 100% boosted | 5705 | 8 | -1 | 17 | 0.09 |
| 50% | 100% boosted | 5462 | 6 | -5 | 16 | 0.26 |
| 60% | 100% boosted | 5547 | 8 | -4 | 19 | 0.18 |
| 70% | 100% boosted | 5607 | 9 | -3 | 21 | 0.14 |
| 80% | 100% boosted | 4737 | -4 | -18 | 10 | 0.52 |

*Note: All tests are comparing the given scenario for proactive vaccination (table rows) against the baseline scenario. Statistically significant results are highlighted by bold text for* p*-values. Colors indicate the direction of change: red indicates that proactive vaccination significantly increased the duration of epidemics. The proportion proactive column gives the proportion of population proactively vaccinated; proportion boosted is the proportion of those vaccinated which received a second, boosting vaccination. MW test statistic is the Mann-Whitney* U *test statistic. The Estimate column gives the Mann-Whitney* U *difference estimate, and the 95% confidence interval is given in the 95% CI (L) (lower) and 95% CI (U) (upper) columns.*

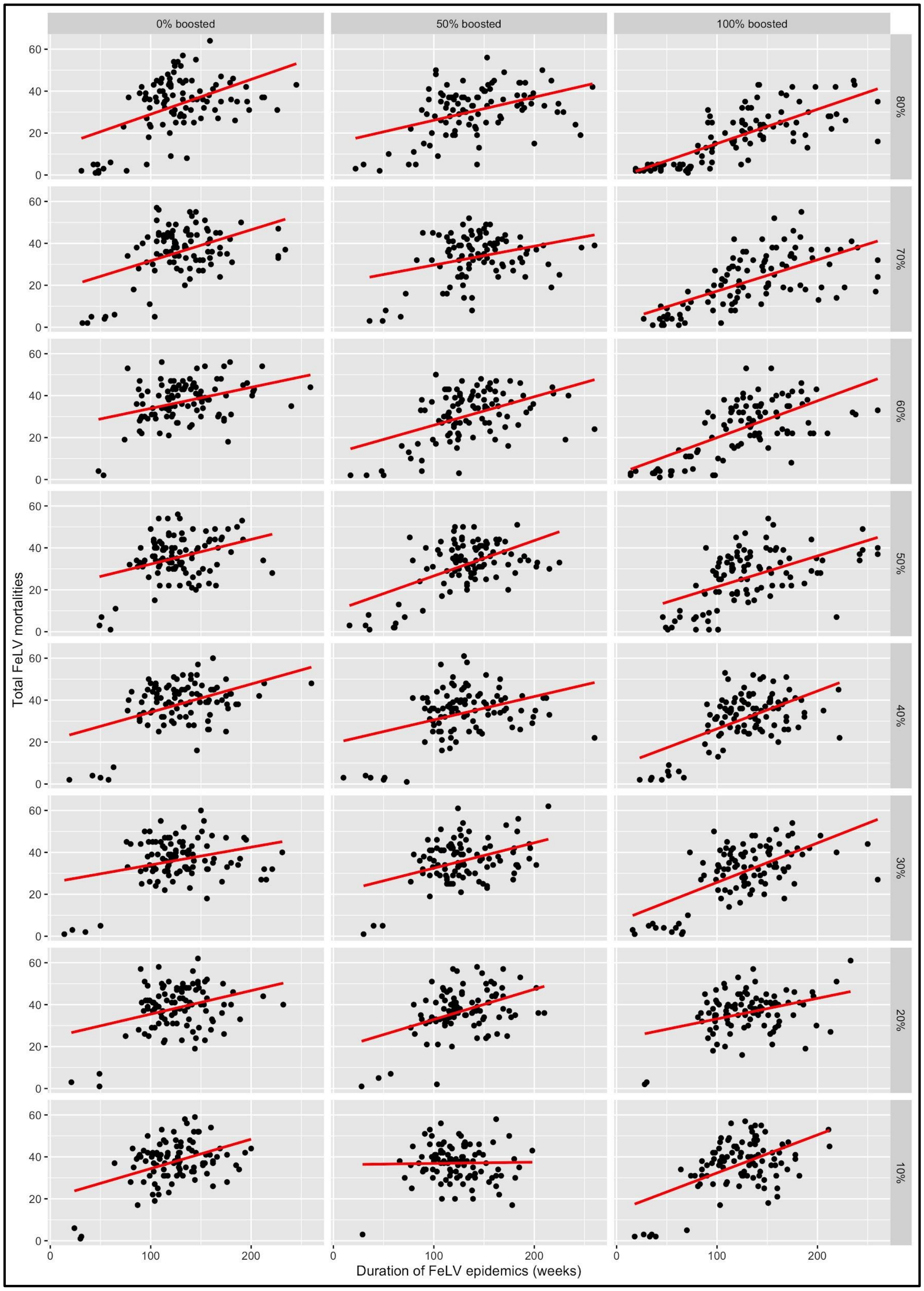

**Figure S4:** Scatterplots of simulated FeLV epidemic durations against mortalities with proactive vaccination alone. Red lines are the fitted linear models for mortalities as a function of epidemic duration. Panel rows represent the proportion of the population proactively vaccinated; columns represent ratios of vaccine efficacy among vaccinates. Vaccination without a booster was assumed to have 40% efficacy, versus 80% efficacy with a boosting inoculation. Each histogram plot represents the results of 100 simulations.

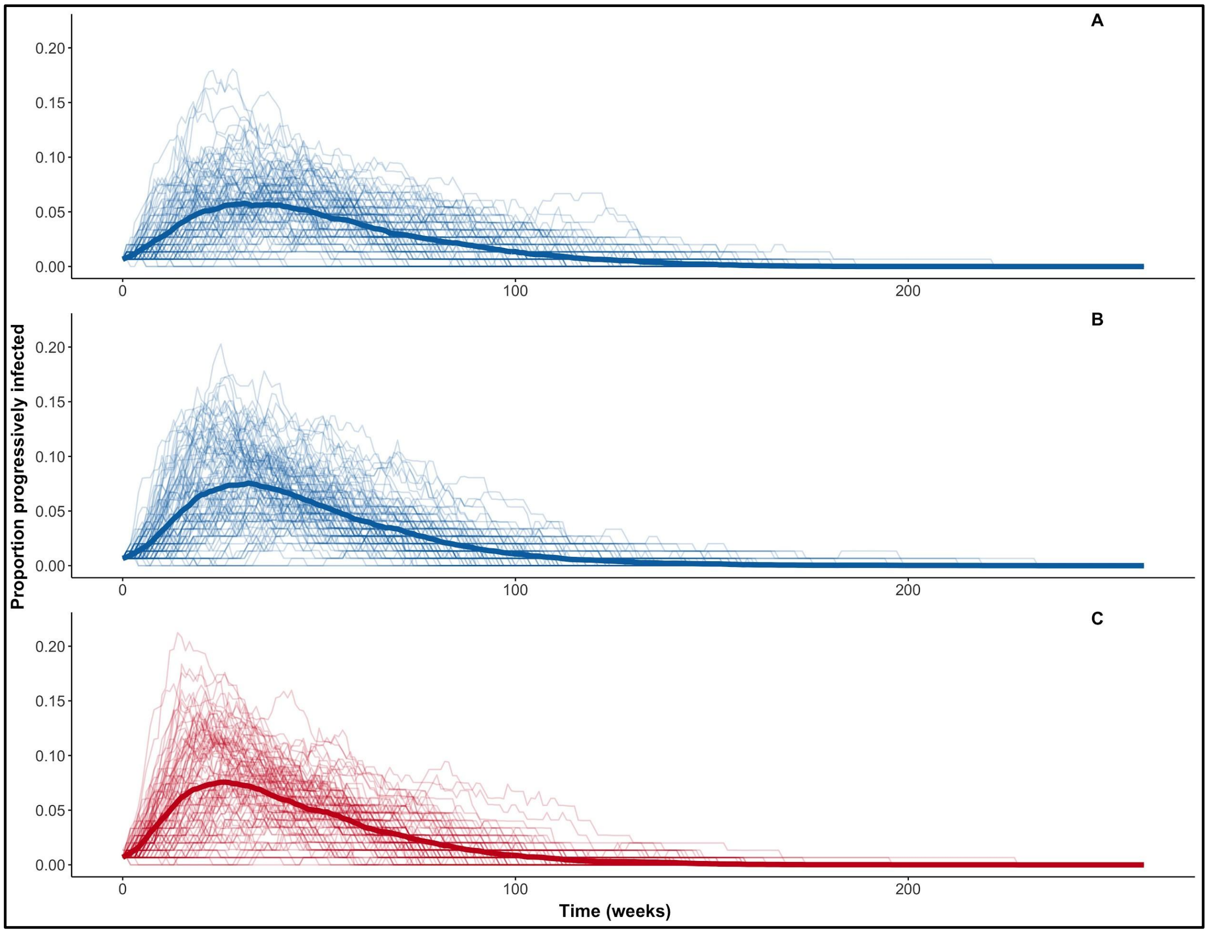

**Figure S5:** Epidemic curves showing the proportion of the population progressively infected over time when (A) 40%, (B) 20%, and (C) 0% of the population was proactively vaccinated, with all vaccinates receiving a booster. Lighter lines show individual simulation results; dark lines show mean values across all simulations. Panels A and B are colored blue to reflect proactive vaccination in use; panel C is colored red to reflect that no interventions are in use.

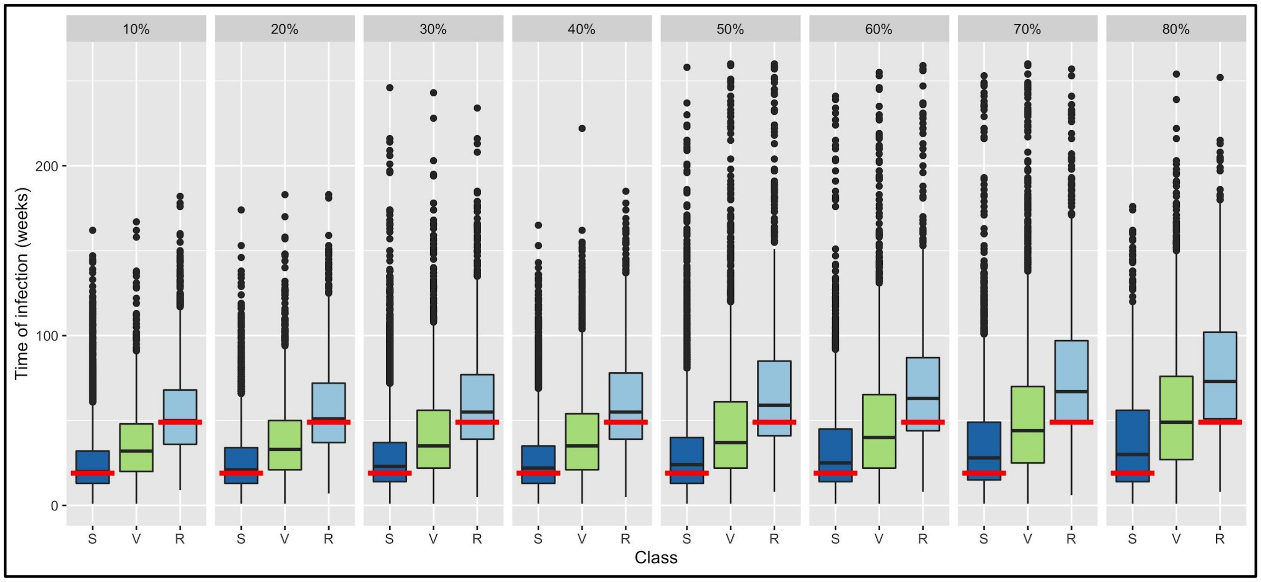

**Figure S6:** Boxplots of the time of infection with proactive vaccination alone. “Class” along the x-axis shows the status of individuals at the time of infection: S = susceptible; V = vaccinated; R = susceptible respawned individual. Panels represent the proportion of the population proactively vaccinated, with all vaccinates receiving a booster. Red horizontal lines in each panel show the median time of infection for susceptibles and susceptible respawns with no interventions (no individuals are vaccinated with no interventions). Results in each panel are given for 100 simulations.

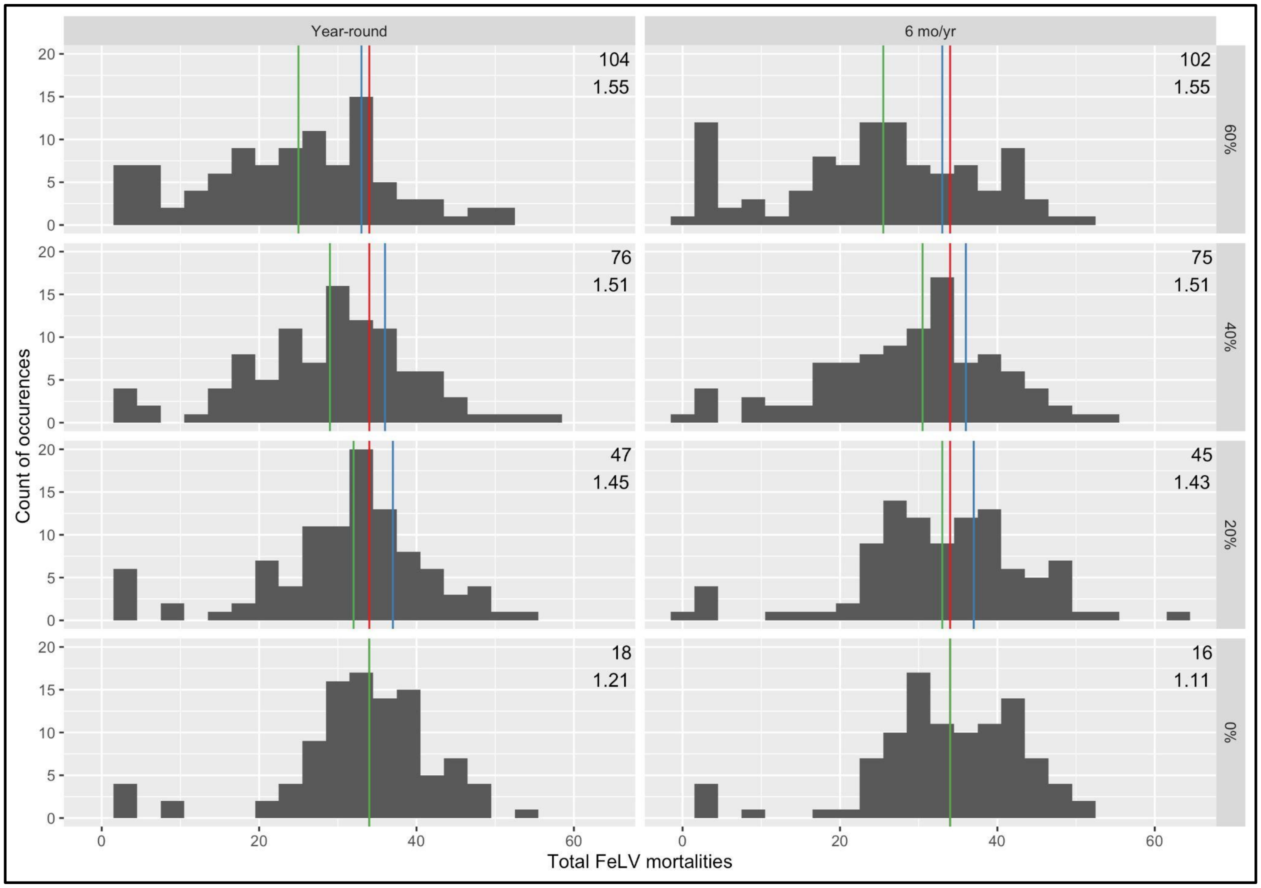

**Figure S7:** Subset of histograms of FeLV mortalities with both proactive and reactive vaccination in the context of vaccination effort. Results shown are for randomly distributed reactive vaccination starting 52 weeks after epidemic onset. Numbers in the upper right of each histogram represent (above) median total number of vaccinated individuals (both proactive and reactive), and (below) median value for the total vaccinations used per total vaccinated individuals. Panel rows represent the proportion of the population proactively vaccinated (with 50% boosted vaccination); note that the bottom row, with 0% proactive vaccination, demonstrates reactive vaccination alone. Columns represent duration of reactive vaccination per year. Red vertical lines indicate the median number of mortalities from simulations without interventions; blue lines for proactive vaccination alone; green lines for the given combination of reactive and proactive vaccination (note green lines overlap red lines in the 0% proactive vaccination scenarios).

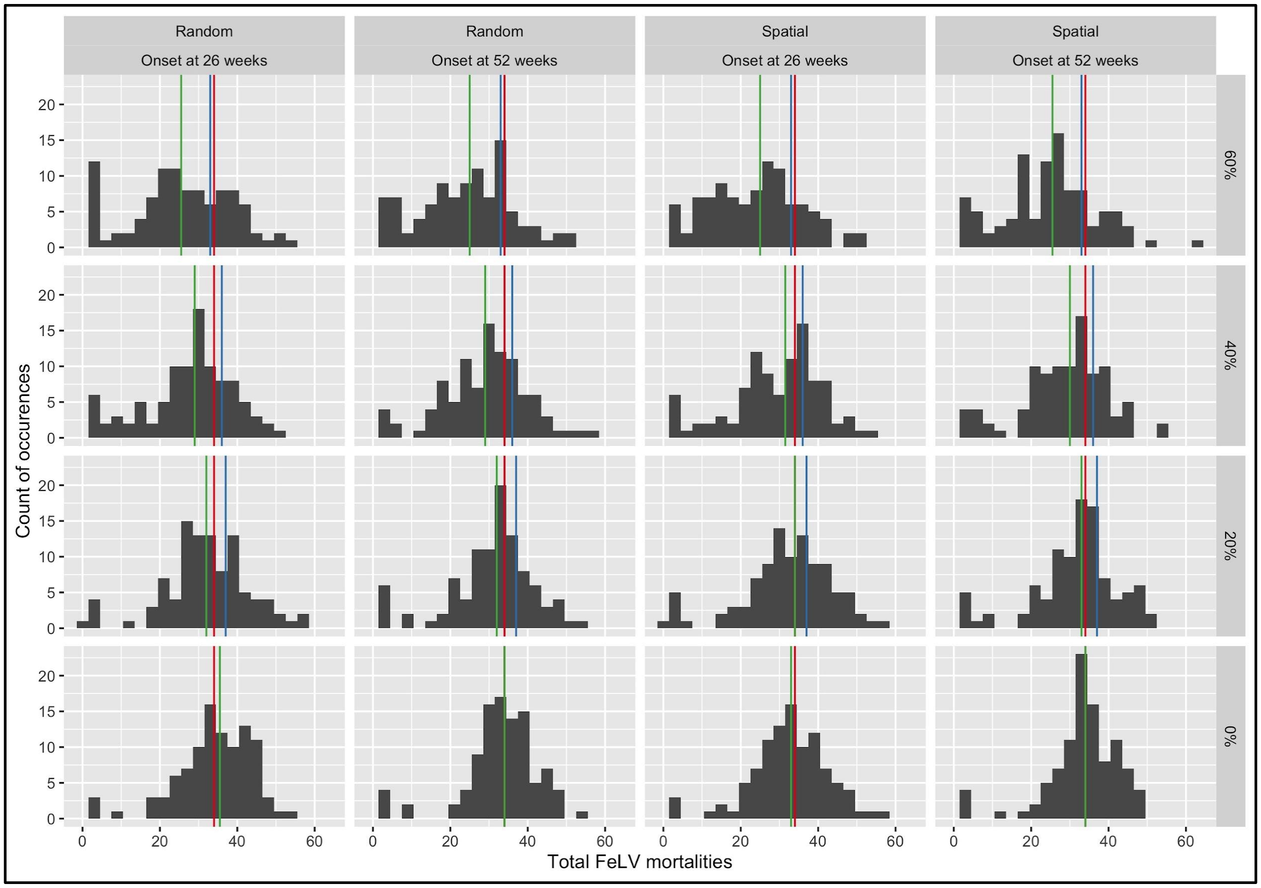

**Figure S8:** Histograms of FeLV mortalities in simulated epidemics with both reactive and proactive vaccination. Results shown are for reactive vaccination performed year-round. Panel rows represent the proportion of the population proactively vaccinated (with 50% of those vaccinates receiving boosted vaccination); columns represent both distribution strategy (random versus spatial) and timing of onset of reactive vaccination after epidemic initiation (26 or 52 weeks). Red vertical lines indicate the median number of mortalities from simulations without interventions; blue lines for proactive vaccination alone; green lines for the given combination of reactive and proactive vaccination. Each histogram plot represents the results of 100 simulations.

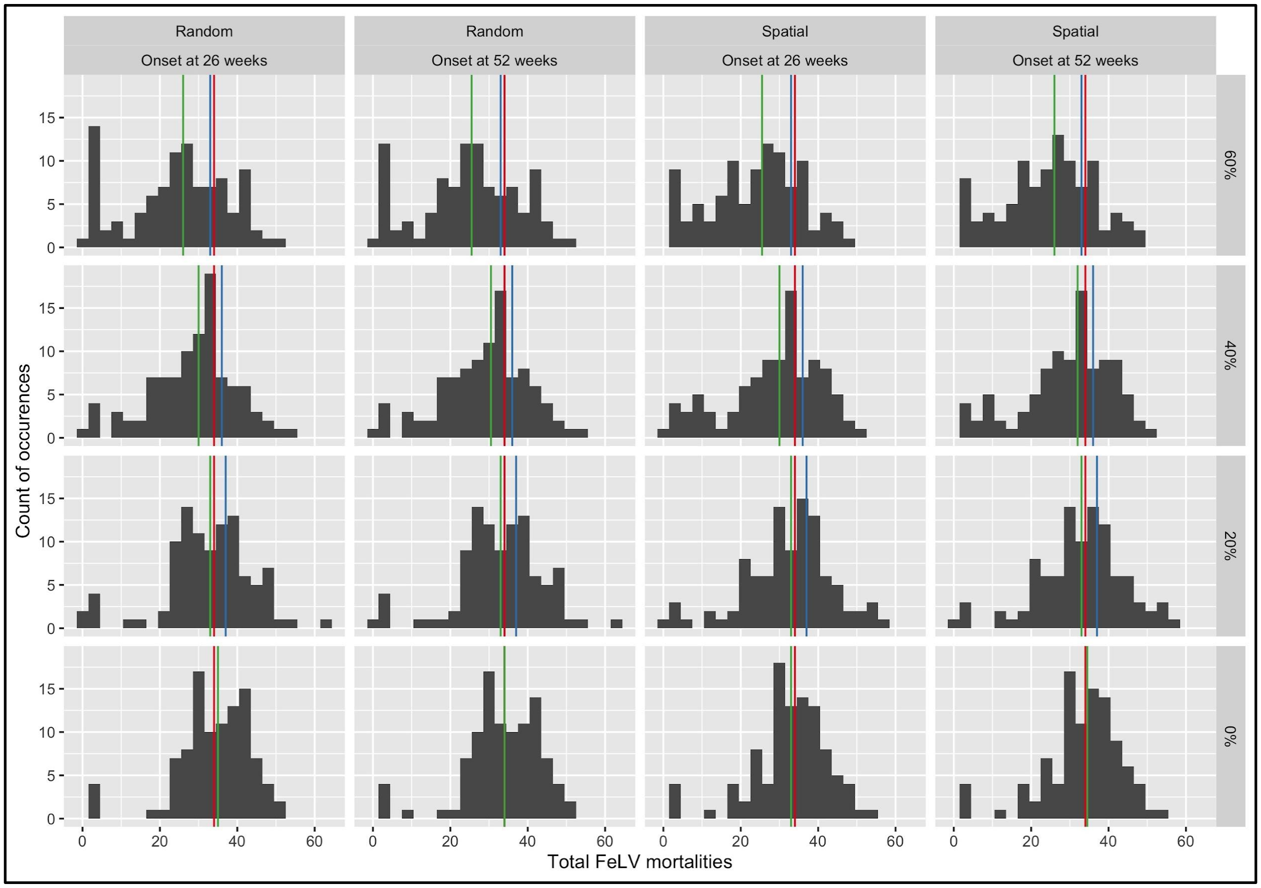

**Figures S9:** Histograms of FeLV mortalities in simulated epidemics with both reactive and proactive vaccination. Results shown are for reactive vaccination performed for six months per year. Panel rows represent the proportion of the population proactively vaccinated (with 50% of those vaccinates receiving boosted vaccination); columns represent both distribution strategy (random versus spatial) and timing of onset of reactive vaccination after epidemic initiation (26 or 52 weeks). Red vertical lines indicate the median number of mortalities from simulations without interventions; blue lines for proactive vaccination alone; green lines for the given combination of reactive and proactive vaccination. Each histogram plot represents the results of 100 simulations.

**Table S4: Mann-Whitney *U* results for mortalities under reactive vaccination**

| Duration of reactive | Distribution strategy | Onset of reactive | Proportion proactive | MW test statistic | Estimate | 95% CI (L) | 95% CI (U) | *p-value* |
|---|---|---|---|---|---|---|---|---|
| 6 months | random | 26 | 0% | 5308 | 1 | -2 | 3 | 0.45 |
| | | 26 | 20% | 4705 | -1 | -4 | 2 | 0.47 |
| | | 26 | 40% | 3722 | -5 | -7 | -2 | **<0.01** |
| | | 26 | 60% | 2998 | -8 | -12 | -5 | **<0.001** |
| | | 52 | 0% | 5116 | 0 | -2 | 3 | 0.78 |
| | | 52 | 20% | 4733 | -1 | -4 | 2 | 0.51 |
| | | 52 | 40% | 3829 | -4 | -7 | -1 | **<0.01** |
| | | 52 | 60% | 3014 | -8 | -12 | -5 | **<0.001** |
| | spatial | 26 | 0% | 4830 | -1 | -3 | 2 | 0.68 |
| | | 26 | 20% | 4708 | -1 | -4 | 2 | 0.48 |
| | | 26 | 40% | 3920 | -4 | -7 | -1 | **<0.01** |
| | | 26 | 60% | 2709 | -9 | -12 | -6 | **<0.001** |
| | | 52 | 0% | 4976 | 0 | -3 | 2 | 0.95 |
| | | 52 | 20% | 4796 | -1 | -4 | 2 | 0.62 |
| | | 52 | 40% | 4075 | -3 | -6 | 0 | **0.02** |
| | | 52 | 60% | 2794 | -9 | -12 | -6 | **<0.001** |
| All year | random | 26 | 0% | 5273 | 1 | -2 | 3 | 0.51 |
| | | 26 | 20% | 4522 | -2 | -4 | 1 | 0.24 |
| | | 26 | 40% | 3479 | -5 | -8 | -3 | **<0.001** |
| | | 26 | 60% | 3210 | -8 | -11 | -4 | **<0.001** |
| | | 52 | 0% | 5000 | 0 | -2 | 2 | 1 |
| | | 52 | 20% | 4366 | -2 | -5 | 1 | 0.12 |
| | | 52 | 40% | 3708 | -5 | -7 | -2 | **<0.01** |

| | | | | | | | | |
|---|---|---|---|---|---|---|---|---|
| | | 52 | 60% | 2752 | -9 | -12 | -6 | **<0.001** |
| | spatial | 26 | 0% | 4744 | -1 | -3 | 2 | 0.53 |
| | | 26 | 20% | 4778 | -1 | -4 | 2 | 0.59 |
| | | 26 | 40% | 4034 | -4 | -6 | -1 | **0.02** |
| | | 26 | 60% | 2691 | -10 | -13 | -7 | **<0.001** |
| | | 52 | 0% | 5074 | 0 | -2 | 3 | 0.86 |
| | | 52 | 20% | 4564 | -1 | -4 | 1 | 0.29 |
| | | 52 | 40% | 3735 | -4 | -7 | -2 | **<0.01** |
| | | 52 | 60% | 2763 | -9 | -12 | -6 | **<0.001** |

*Note: All tests are comparing the given scenario for reactive vaccination (table rows) against the baseline scenario. Statistically significant results are highlighted by bold text for* p*-values. Color indicates the direction of change: blue indicates that the given reactive vaccination scenario significantly reduced mortalities. The duration reactive gives the annual duration of reactive vaccination; distribution strategy indicates if reactive vaccinations were distributed randomly or spatially in an attempted vaccine barrier. Onset of reactive is the time (in weeks) after epidemic initiation at which reactive vaccination began; the proportion proactive column gives the proportion of population proactively vaccinated prior to an epidemic. MW test statistic is the Mann-Whitney* U *test statistic. The Estimate column gives the Mann-Whitney* U *difference estimate, and the 95% confidence interval is given in the 95% CI (L) (lower) and 95% CI (U) (upper) columns.*

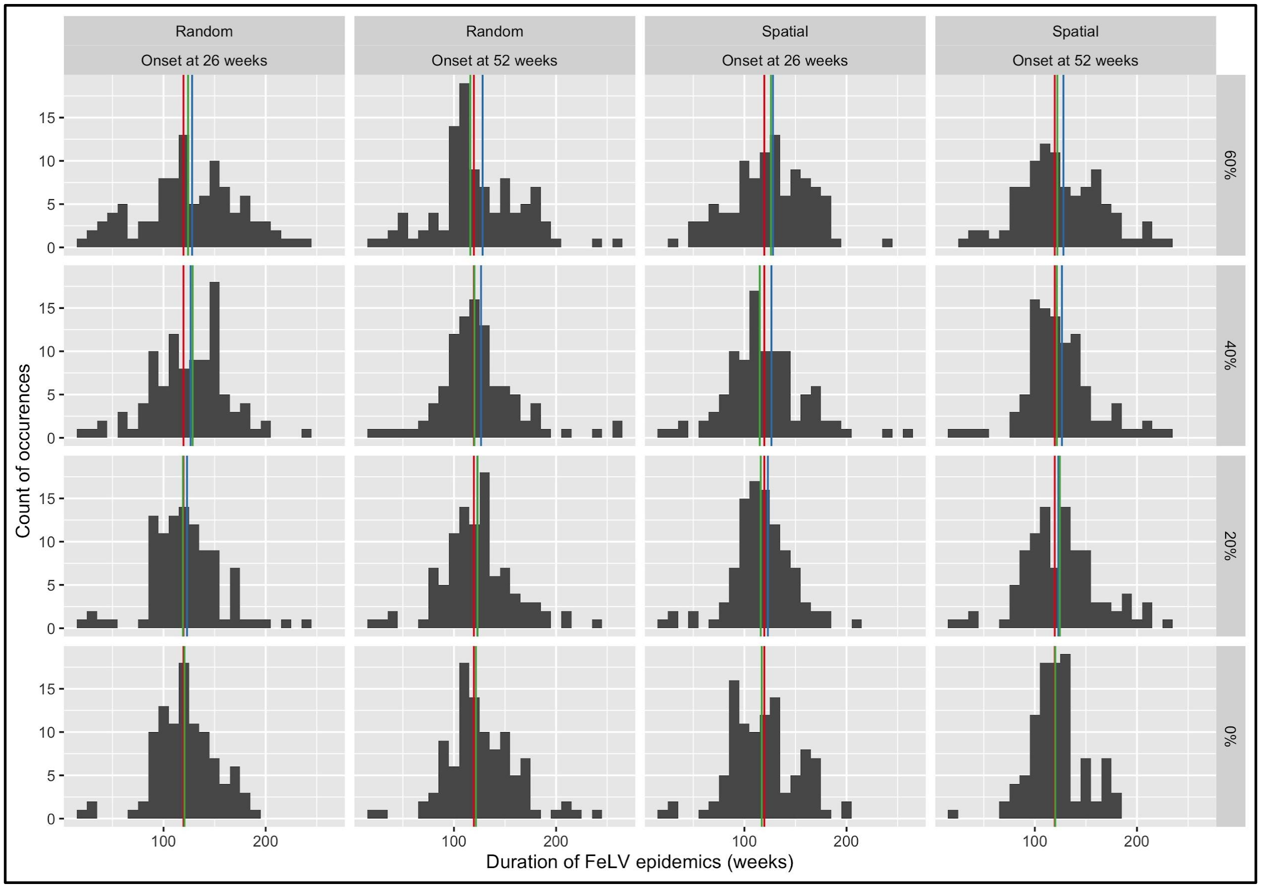

**Figure S10:** Histograms of the duration of simulated FeLV epidemics with both reactive and proactive vaccination. Results shown are for reactive vaccination performed year-round. Panel rows represent the proportion of the population proactively vaccinated (with 50% of those vaccinates receiving boosted vaccination); columns represent both distribution strategy (random versus spatial) and timing of onset of reactive vaccination after epidemic initiation (26 or 52 weeks). Red vertical lines indicate the median duration without interventions; blue lines for proactive vaccination alone; green lines for the given combination of reactive and proactive vaccination. Each histogram plot represents the results of 100 simulations.

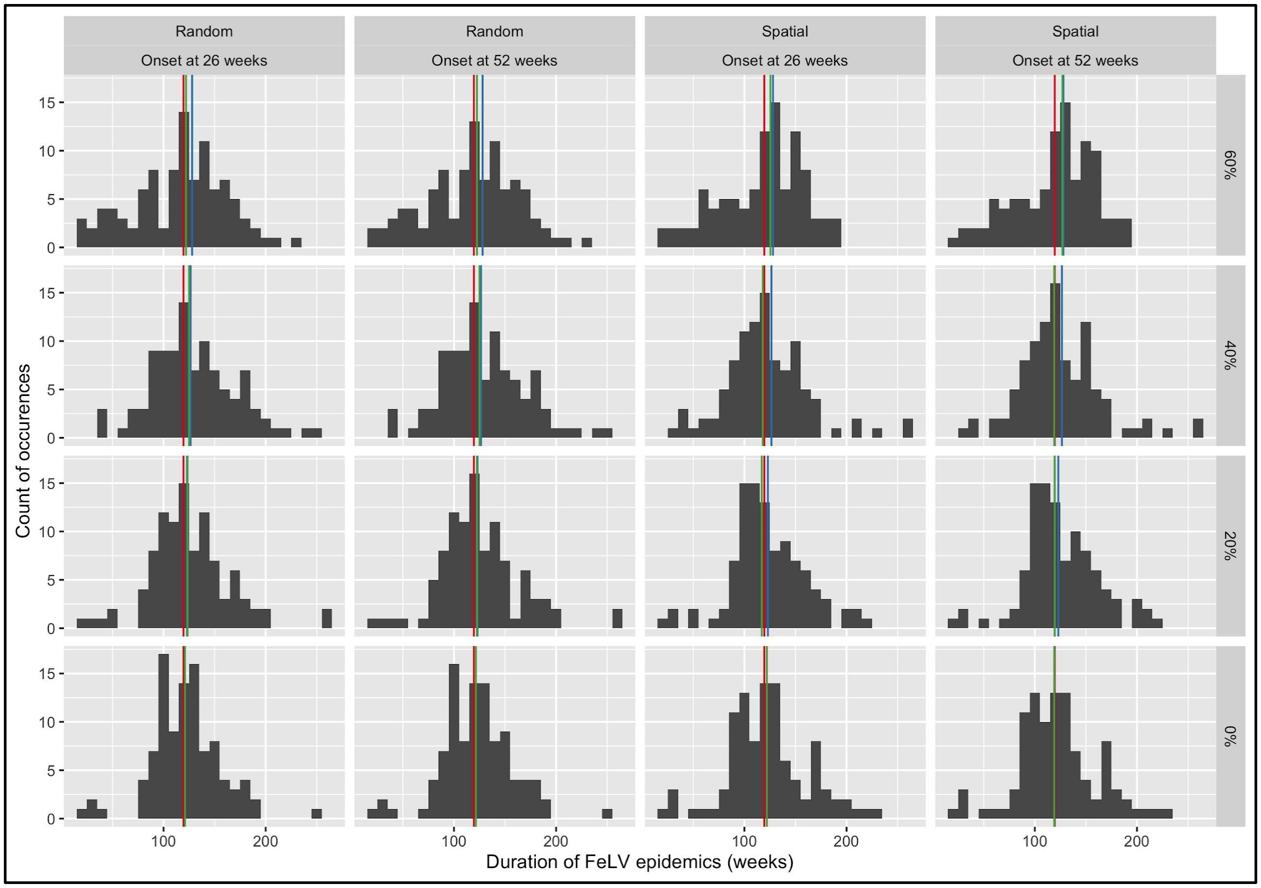

**Figure S11:** Histograms of the duration of simulated FeLV epidemics with both reactive and proactive vaccination. Results shown are for reactive vaccination performed for six months per year. Panel rows represent the proportion of the population proactively vaccinated (with 50% of those vaccinates receiving boosted vaccination); columns represent both distribution strategy (random versus spatial) and timing of onset of reactive vaccination after epidemic initiation (26 or 52 weeks). Red vertical lines indicate the median duration without interventions; blue lines for proactive vaccination alone; green lines for the given combination of reactive and proactive vaccination. Each histogram plot represents the results of 100 simulations.

**Table S5: Mann-Whitney *U* results for epidemic durations under reactive vaccination**

| Duration of reactive | Distribution strategy | Onset of reactive | Proportion proactive | MW test statistic | Estimate | 95% CI (L) | 95% CI (U) | *p-value* |
|---|---|---|---|---|---|---|---|---|
| 6 months | random | 26 | 0% | 5114 | 1 | -7 | 10 | 0.78 |
| | | 26 | 20% | 5292 | 3 | -5 | 13 | 0.48 |
| | | 26 | 40% | 5607 | 8 | -3 | 18 | 0.14 |
| | | 26 | 60% | 4869 | -2 | -13 | 9 | 0.75 |
| | | 52 | 0% | 5175 | 2 | -7 | 10 | 0.67 |
| | | 52 | 20% | 5208 | 2 | -6 | 12 | 0.61 |
| | | 52 | 40% | 5597 | 7 | -2 | 17 | 0.15 |
| | | 52 | 60% | 4941 | -1 | -12 | 10 | 0.89 |
| | spatial | 26 | 0% | 5065 | 1 | -9 | 10 | 0.87 |
| | | 26 | 20% | 5152 | 1 | -7 | 10 | 0.71 |
| | | 26 | 40% | 4905 | -1 | -10 | 8 | 0.82 |
| | | 26 | 60% | 5062 | 1 | -10 | 11 | 0.88 |
| | | 52 | 0% | 5006 | 0 | -9 | 10 | 0.99 |
| | | 52 | 20% | 5324 | 3 | -5 | 12 | 0.43 |
| | | 52 | 40% | 5134 | 2 | -7 | 10 | 0.74 |
| | | 52 | 60% | 5229 | 3 | -8 | 13 | 0.58 |
| All year | random | 26 | 0% | 5123 | 1 | -7 | 10 | 0.77 |
| | | 26 | 20% | 5000 | 0 | -8 | 9 | 1 |
| | | 26 | 40% | 5409 | 5 | -5 | 14 | 0.32 |
| | | 26 | 60% | 5476 | 7 | -5 | 17 | 0.25 |
| | | 52 | 0% | 5353 | 4 | -5 | 12 | 0.39 |
| | | 52 | 20% | 5035 | 0 | -9 | 9 | 0.93 |
| | | 52 | 40% | 4998 | 0 | -9 | 9 | 1 |

| | | | | | | | | |
|---|---|---|---|---|---|---|---|---|
| | | 52 | 60% | 5082 | 1 | -9 | 11 | 0.84 |
| | spatial | 26 | 0% | 4719 | -3 | -12 | 6 | 0.49 |
| | | 26 | 20% | 4682 | -3 | -11 | 5 | 0.44 |
| | | 26 | 40% | 4723 | -3 | -12 | 6 | 0.5 |
| | | 26 | 60% | 5402 | 5 | -5 | 15 | 0.33 |
| | | 52 | 0% | 4992 | 0 | -9 | 8 | 0.98 |
| | | 52 | 20% | 5228 | 2 | -7 | 12 | 0.58 |
| | | 52 | 40% | 5085 | 1 | -7 | 10 | 0.84 |
| | | 52 | 60% | 5261 | 3 | -7 | 13 | 0.53 |

*Note: All tests are comparing the given scenario for reactive vaccination (table rows) against the baseline scenario. No statistically significant results were found. The duration reactive gives the annual duration of reactive vaccination; distribution strategy indicates if reactive vaccinations were distributed randomly or spatially in an attempted vaccine barrier. Onset of reactive is the time (in weeks) after epidemic initiation at which reactive vaccination began; the proportion proactive column gives the proportion of population proactively vaccinated prior to an epidemic. MW test statistic is the Mann-Whitney* U *test statistic. The Estimate column gives the Mann-Whitney* U *difference estimate, and the 95% confidence interval is given in the 95% CI (L) (lower) and 95% CI (U) (upper) columns.*

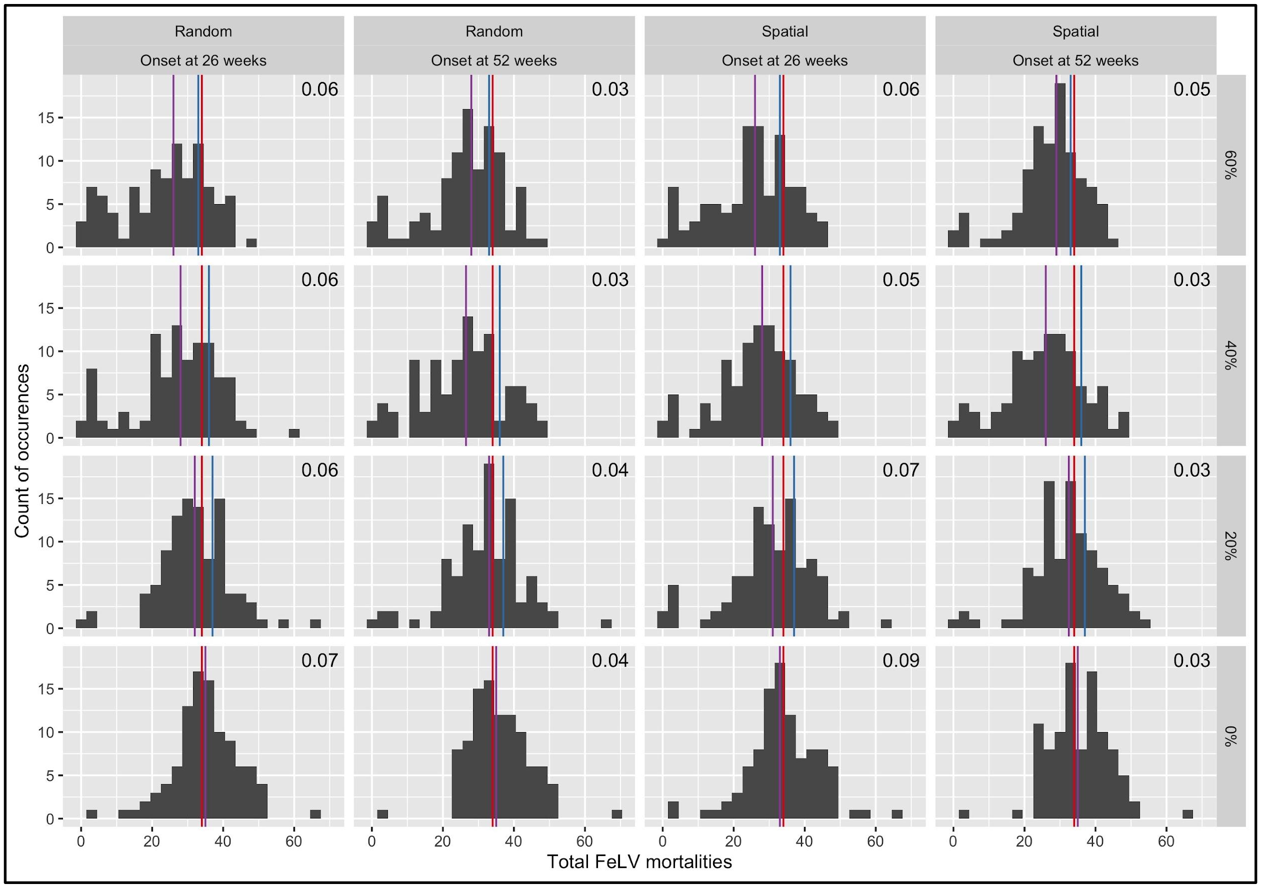

**Figure S12:** Histograms of FeLV mortalities in simulated epidemics with both reactive test-and-removal and proactive vaccination. Panel rows represent the proportion of the population proactively vaccinated; columns represent both capture/testing strategy (random versus spatial) and timing of onset of reactive test-and-removal after epidemic initiation (26 or 52 weeks). Red vertical lines indicate the median number of mortalities from simulations without interventions; blue lines for proactive vaccination alone; purple lines for the given combination of reactive test-and-removal and proactive vaccination. Numbers in the upper right of each histogram represent the median value for the proportion of capture events that resulted in a removal or humane euthanasia. Each histogram plot represents the results of 100 simulations.

**Table S6: Mann-Whitney *U* results for mortalities under reactive test-and-removal**

| Capture/Testing Strategy | Onset of reactive | Proportion proactive | MW test statistic | Estimate | 95% CI (L) | 95% CI (U) | *p-value* |
|---|---|---|---|---|---|---|---|
| random | 26 | 0% | 5403 | 1 | -1 | 4 | 0.33 |
| | 26 | 20% | 4435 | -2 | -4 | 1 | 0.17 |
| | 26 | 40% | 3430 | -6 | -9 | -3 | **<0.001** |
| | 26 | 60% | 2707 | -9 | -13 | -6 | **<0.001** |
| | 52 | 0% | 5472 | 1 | -1 | 4 | 0.25 |
| | 52 | 20% | 4519 | -2 | -4 | 1 | 0.24 |
| | 52 | 40% | 3085 | -8 | -11 | -5 | **<0.001** |
| | 52 | 60% | 3196 | -6 | -9 | -4 | **<0.001** |
| spatial | 26 | 0% | 5078 | 0 | -2 | 3 | 0.85 |
| | 26 | 20% | 4283 | -2 | -5 | 0 | 0.08 |
| | 26 | 40% | 3260 | -6 | -9 | -3 | **<0.001** |
| | 26 | 60% | 2975 | -8 | -11 | -5 | **<0.001** |
| | 52 | 0% | 5433 | 1 | -1 | 4 | 0.29 |
| | 52 | 20% | 4548 | -2 | -4 | 1 | 0.27 |
| | 52 | 40% | 2994 | -8 | -11 | -5 | **<0.001** |
| | 52 | 60% | 3198 | -6 | -8 | -3 | **<0.001** |

*Note: All tests are comparing the given scenario for reactive test-and-removal (table rows) against the baseline scenario. Statistically significant results are highlighted by bold text for* p-*values. Color indicates the direction of change: blue indicates that the given reactive test-and-removal scenario significantly reduced mortalities. The capture/testing strategy column indicates if captures for reactive test-and-removal were random or spatially targeted. Onset of reactive is the time (in weeks) after epidemic initiation at which reactive test-and-removal began; the proportion proactive column gives the proportion of population proactively vaccinated prior to an* *epidemic. MW test statistic is the Mann-Whitney* U *test statistic. The Estimate column gives the Mann-Whitney* U *difference estimate, and the 95% confidence interval is given in the 95% CI (L) (lower) and 95% CI (U) (upper) columns.*

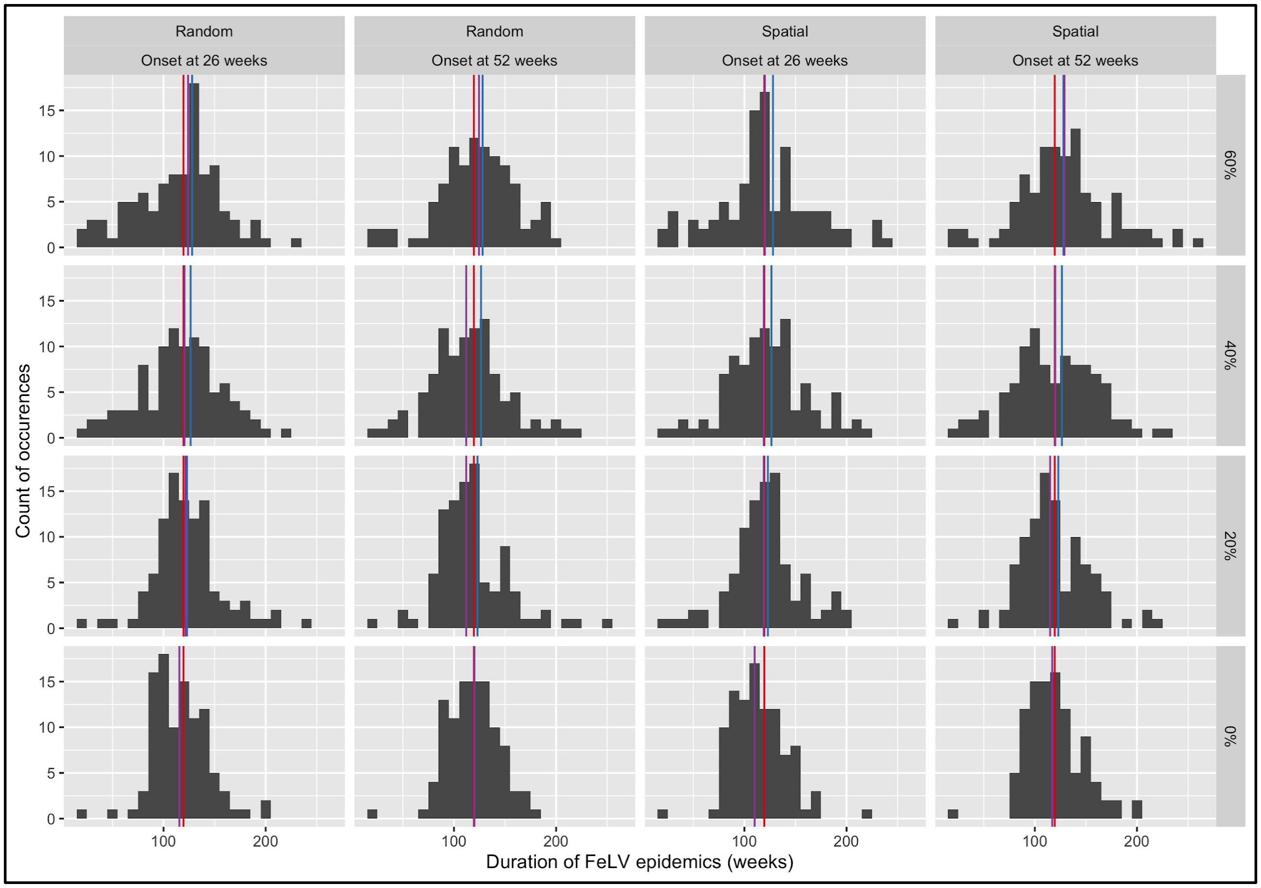

**Figure S13:** Histograms of the duration of simulated FeLV epidemics with both reactive test-and-removal and proactive vaccination. Panel rows represent the proportion of the population proactively vaccinated; columns represent both capture/testing strategy (random versus spatial) and timing of onset of reactive test-and-removal after epidemic initiation (26 or 52 weeks). Red vertical lines indicate the median duration without interventions; blue lines for proactive vaccination alone; purple lines for the given combination of reactive test-and-removal and proactive vaccination. Each histogram plot represents the results of 100 simulations.

**Table S7: Mann-Whitney *U* results for epidemic durations under reactive test-and-removal**

| Capture/Testing Strategy | Onset of reactive | Proportion proactive | MW test statistic | Estimate | 95% CI (L) | 95% CI (U) | *p-value* |
|---|---|---|---|---|---|---|---|
| random | 26 | 0% | 4425 | -5 | -13 | 2 | 0.16 |
| | 26 | 20% | 5053 | 0 | -8 | 9 | 0.9 |
| | 26 | 40% | 4766 | -3 | -13 | 7 | 0.57 |
| | 26 | 60% | 4745 | -4 | -14 | 7 | 0.53 |
| | 52 | 0% | 4765 | -2.5 | -10 | 6 | 0.57 |
| | 52 | 20% | 4493 | -5 | -14 | 3 | 0.22 |
| | 52 | 40% | 4275 | -8 | -17 | 1 | 0.08 |
| | 52 | 60% | 5228 | 3 | -7 | 12 | 0.58 |
| spatial | 26 | 0% | 4158 | -8 | -16 | 0 | **0.04** |
| | 26 | 20% | 4886 | -1 | -10 | 8 | 0.78 |
| | 26 | 40% | 4888 | -1 | -10 | 9 | 0.79 |
| | 26 | 60% | 5010 | 0 | -10 | 10 | 0.98 |
| | 52 | 0% | 4668 | -3 | -11 | 5 | 0.42 |
| | 52 | 20% | 4763 | -2 | -11 | 6 | 0.56 |
| | 52 | 40% | 4784 | -2 | -13 | 8 | 0.6 |
| | 52 | 60% | 5447 | 5.8 | -4 | 16 | 0.28 |

*Note: All tests are comparing the given scenario for reactive test-and-removal (table rows) against the baseline scenario. Statistically significant results are highlighted by bold text for* p-*values. Color indicates the direction of change: blue indicates that the given reactive test-and-removal scenario significantly reduced epidemic duration. The capture/testing strategy column indicates if captures for reactive test-and-removal were random or spatially targeted. Onset of reactive is the time (in weeks) after epidemic initiation at which reactive test-and-removal began;*

*the proportion proactive column gives the proportion of population proactively vaccinated prior to an epidemic. MW test statistic is the Mann-Whitney* U *test statistic. The Estimate column gives the Mann-Whitney* U *difference estimate, and the 95% confidence interval is given in the 95% CI (L) (lower) and 95% CI (U) (upper) columns.*

*Reactive underpass closures*

Reactive underpass closures were ineffective in the absence of proactive vaccination (Figure S14, Table S8). The greatest impact of underpass closures for reducing FeLV mortalities occurred when onset of closure was early (26 weeks after epidemic initiation), lasted for at least 13 weeks, and occurred in conjunction with at least 40-60% of the population being proactively vaccinated (though the clearest effects occurred when at least 60% were proactively vaccinated). Under these conditions, underpass closures synergistically reduced mortalities from baseline scenarios (as few as a median of 22 mortalities with underpass closures) and increased the probability that an epidemic would fail (maximum of 75 failed epidemics per 100 successes with underpass closures; Figure S17). In some cases, simulations showed a marked decrease in transmission during the period of underpass closures, but reopening often resulted in a subsequent resurgence of infections (Figure S15). Underpass closures had no clear impact on the duration of outbreaks (Figure S16, Table S9).

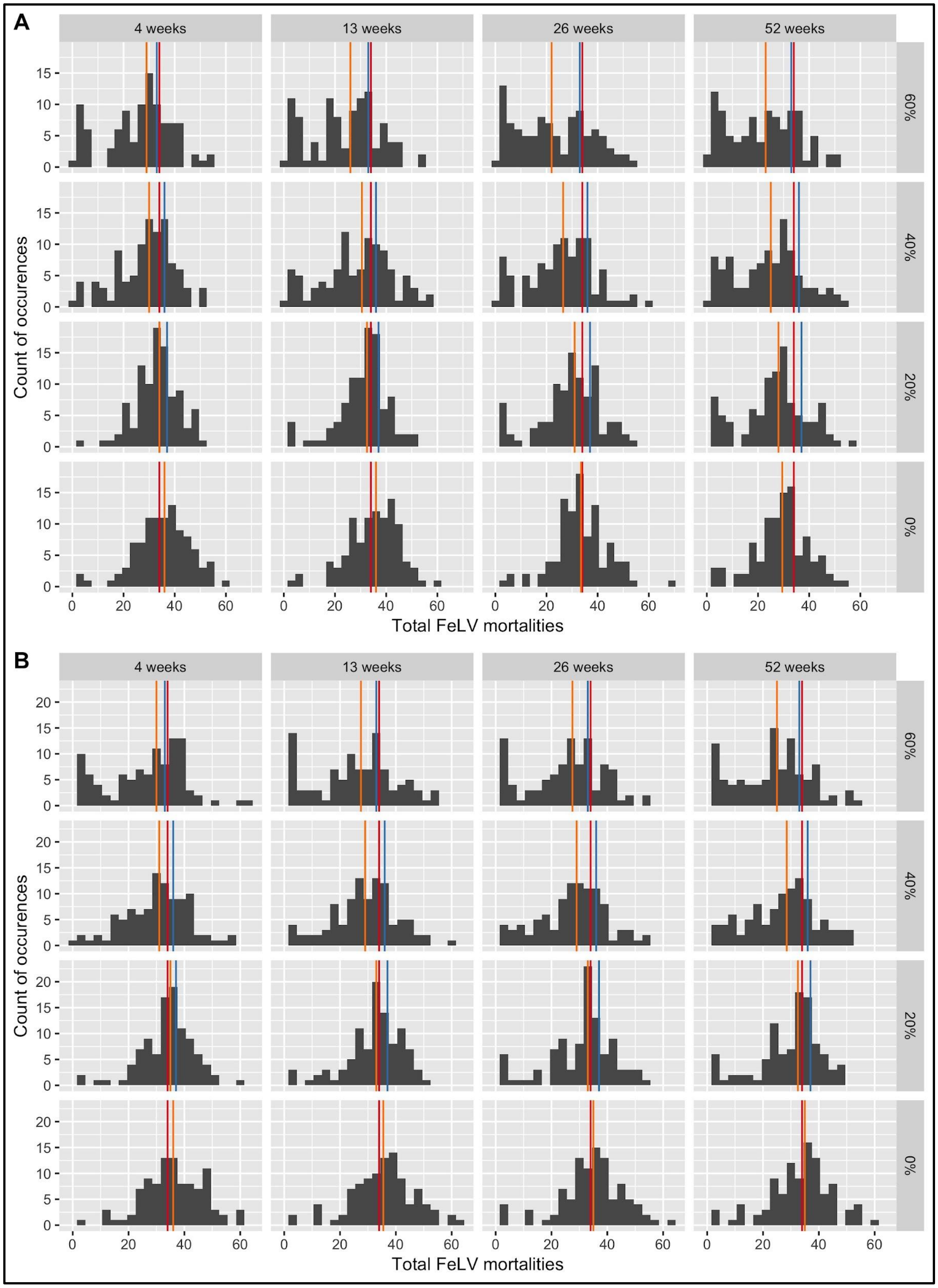

**Figure S14:** Histograms of FeLV mortalities in simulated epidemics with both reactive underpass closures and proactive vaccination. Plot subsections represent (A) onset of underpass closures 26 weeks after epidemic initiation, and (B) onset 52 weeks after initiation. Panel rows represent the proportion of the population proactively vaccinated; columns show the duration of underpass closures. Red vertical lines indicate the median number of mortalities from simulations without interventions; blue lines for proactive vaccination alone; orange lines for the given combination of reactive underpass closure and proactive vaccination. Each histogram plot represents the results of 100 simulations.

**Table S8: Mann-Whitney *U* results for mortalites under reactive underpass closures**

| Duration of closure | Onset of reactive | Proportion proactive | MW test statistic | Estimate | 95% CI (L) | 95% CI (U) | *p-value* |
|---|---|---|---|---|---|---|---|
| 4 | 26 | 0% | 5509 | 2 | -1 | 5 | 0.21 |
| | 26 | 20% | 4796 | -1 | -3 | 2 | 0.62 |
| | 26 | 40% | 3781 | -4 | -7 | -1 | **<0.01** |
| | 26 | 60% | 3351 | -7 | -10 | -3 | **<0.001** |
| | 52 | 0% | 5669 | 2 | 0 | 5 | 0.1 |
| | 52 | 20% | 5129 | 0 | -2 | 3 | 0.75 |
| | 52 | 40% | 4185 | -3 | -6 | 0 | **0.05** |
| | 52 | 60% | 3572 | -6 | -9 | -3 | **<0.001** |
| 13 | 26 | 0% | 5489 | 2 | -1 | 4 | 0.23 |
| | 26 | 20% | 4327 | -2 | -5 | 0 | 0.1 |
| | 26 | 40% | 3989 | -4 | -8 | -1 | **0.01** |
| | 26 | 60% | 2992 | -9 | -13 | -5 | **<0.001** |
| | 52 | 0% | 5559 | 2 | -1 | 5 | 0.17 |
| | 52 | 20% | 4777 | -1 | -3 | 2 | 0.59 |
| | 52 | 40% | 3728 | -5 | -7 | -2 | **<0.01** |
| | 52 | 60% | 3403 | -7 | -11 | -4 | **<0.001** |

| | | | | | | | |
|---|---|---|---|---|---|---|---|
| 26 | 26 | 0% | 4875 | 0 | -3 | 2 | 0.76 |
| | 26 | 20% | 4027 | -4 | -6 | -1 | **0.02** |
| | 26 | 40% | 3165 | -7 | -11 | -4 | **<0.001** |
| | 26 | 60% | 2909 | -11 | -15 | -7 | **<0.001** |
| | 52 | 0% | 5323 | 1 | -2 | 4 | 0.43 |
| | 52 | 20% | 4504 | -2 | -5 | 1 | 0.23 |
| | 52 | 40% | 3550 | -5 | -8 | -2 | **<0.001** |
| | 52 | 60% | 3220 | -7 | -11 | -4 | **<0.001** |
| 52 | 26 | 0% | 3759 | -4 | -7 | -2 | **<0.01** |
| | 26 | 20% | 3354 | -6 | -10 | -3 | **<0.001** |
| | 26 | 40% | 2898 | -9 | -13 | -6 | **<0.001** |
| | 26 | 60% | 2514 | -12 | -16 | -8 | **<0.001** |
| | 52 | 0% | 4992 | 0 | -3 | 3 | 0.98 |
| | 52 | 20% | 4160 | -3 | -6 | 0 | **0.04** |
| | 52 | 40% | 3403 | -6 | -10 | -3 | **<0.001** |
| | 52 | 60% | 2811 | -9 | -13 | -6 | **<0.001** |

*Note: All tests are comparing the given scenario for reactive underpass closures (table rows) against the baseline scenario. Statistically significant results are highlighted by bold text for* p-*values. Color indicates the direction of change: blue indicates that the given reactive underpass closure scenario significantly reduced mortalities. The duration of closure column indicates the duration (in weeks) that simulated underpasses were closed. Onset of reactive is the time (in weeks) after epidemic initiation at which reactive closures began; the proportion proactive column gives the proportion of population proactively vaccinated prior to an epidemic. MW test statistic is the Mann-Whitney* U *test statistic. The Estimate column gives the Mann-Whitney* U *difference estimate, and the 95% confidence interval is given in the 95% CI (L) (lower) and 95% CI (U) (upper) columns.*

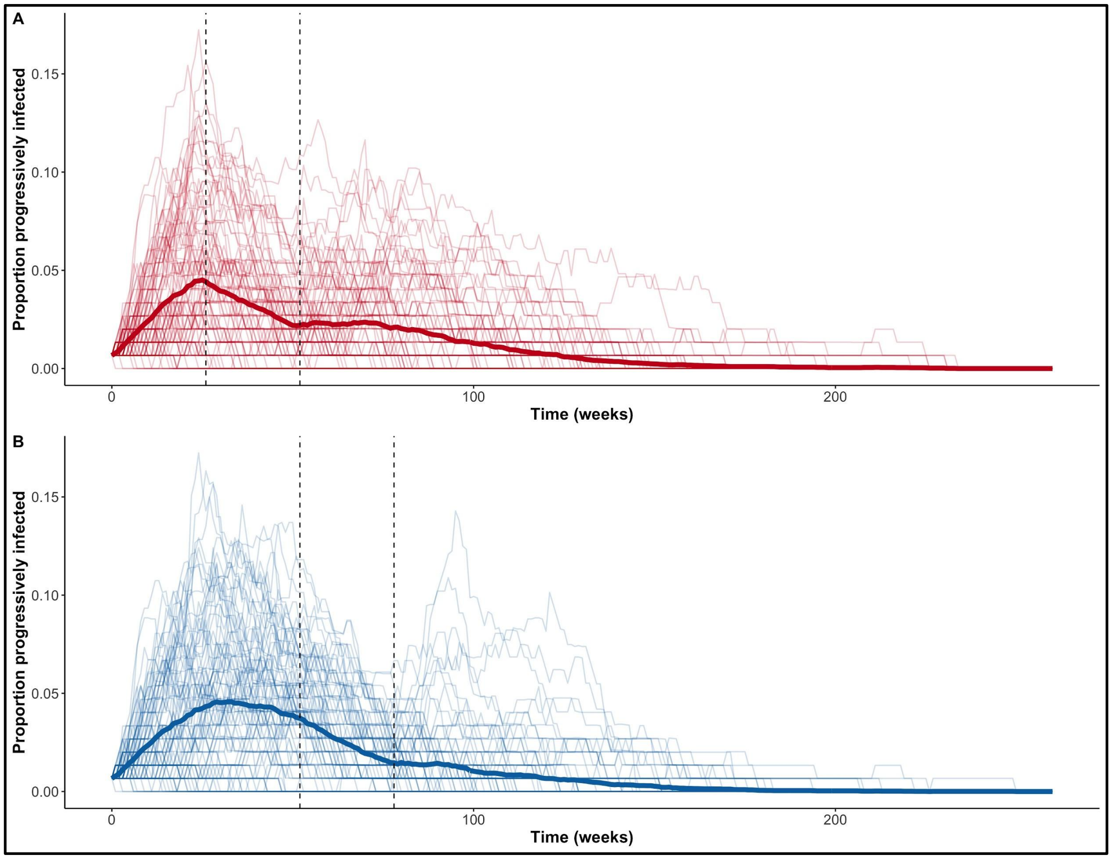

**Figure S15:** Epidemic curves showing the proportion of the population progressively infected over time with underpass closures when 60% of the population was proactively vaccinated. Lighter lines show individual simulation results; dark lines show mean values across all simulations. Vertical black dashed lines show when underpasses were closed (left) and reopened (right). Panel A shows underpasses closed for 26 weeks starting 26 weeks after an epidemic started; panel B shows underpasses closed for 26 weeks starting at 52 weeks.

451

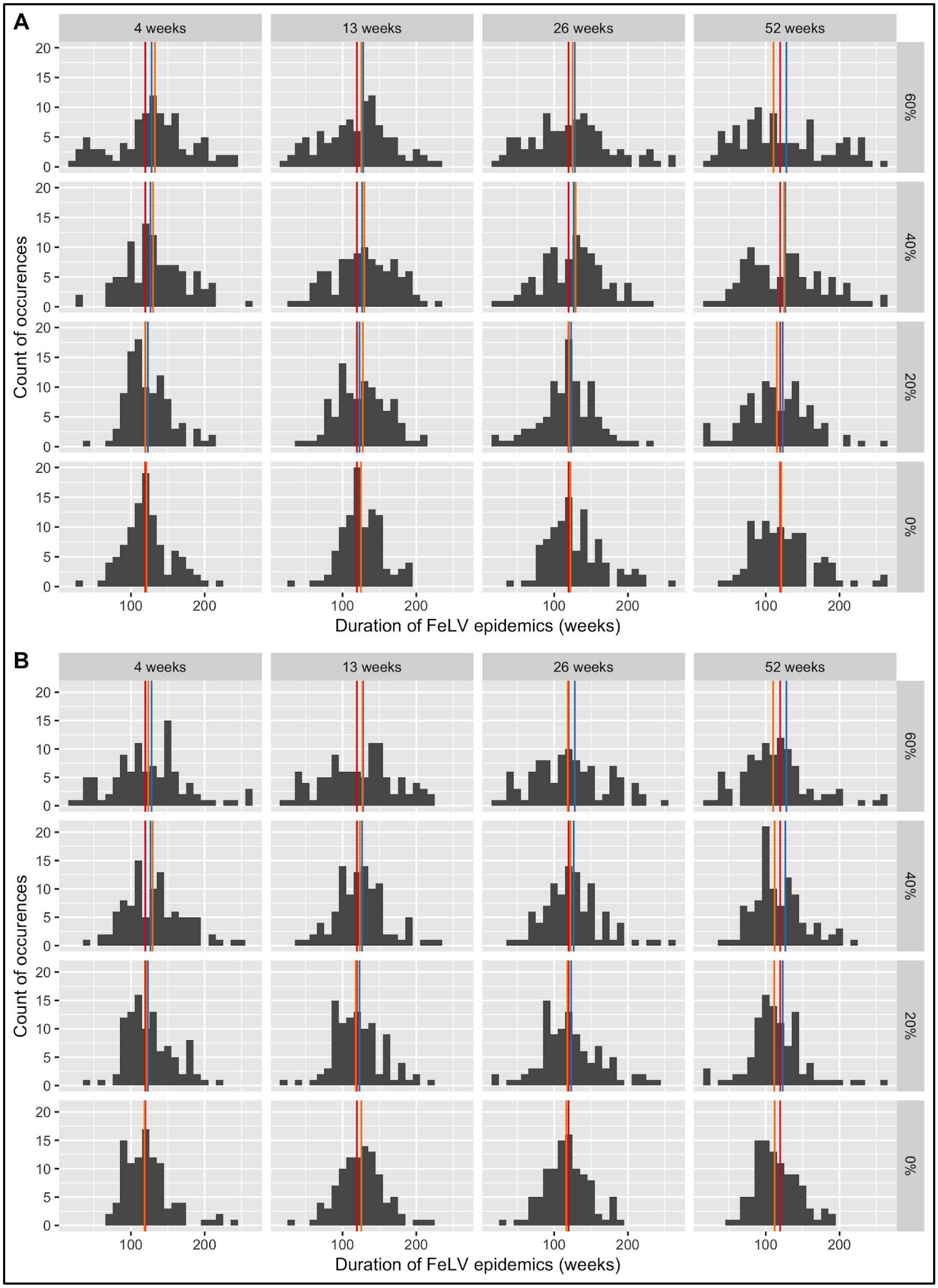
A
4 weeks
13 weeks
26 weeks
52 weeks
60%
40%
20%
0%
Count of occurences
Duration of FeLV epidemics (weeks)
B
4 weeks
13 weeks
26 weeks
52 weeks
60%
40%
20%
0%
Count of occurences
Duration of FeLV epidemics (weeks)

**Figure S16:** Histograms of the duration of simulated FeLV epidemics with both reactive underpass closures and proactive vaccination. Plot subsections represent (A) onset of underpass closures 26 weeks after epidemic initiation, and (B) onset 52 weeks after initiation. Panel rows represent the proportion of the population proactively vaccinated; columns show the duration of underpass closures. Red vertical lines indicate the median duration without interventions; blue lines for proactive vaccination alone; orange lines for the given combination of reactive underpass closure and proactive vaccination. Each histogram plot represents the results of 100 simulations.

**Table S9: Mann-Whitney *U* results for epidemic durations under reactive underpass closures**

| Duration of closure | Onset of reactive | Proportion proactive | MW test statistic | Estimate | 95% CI (L) | 95% CI (U) | *p-value* |
|---|---|---|---|---|---|---|---|
| 4 | 26 | 0% | 5026 | 0 | -9 | 9 | 0.95 |
| | 26 | 20% | 5110 | 1 | -7 | 9 | 0.79 |
| | 26 | 40% | 5849 | 11 | 0 | 20 | **0.04** |
| | 26 | 60% | 5791 | 11 | 0 | 23 | **0.05** |
| | 52 | 0% | 4855 | -2 | -10 | 7 | 0.72 |
| | 52 | 20% | 5267 | 3 | -6 | 12 | 0.52 |
| | 52 | 40% | 5666 | 8 | -2 | 18 | 0.1 |
| | 52 | 60% | 5158 | 2 | -9 | 12 | 0.7 |
| 13 | 26 | 0% | 5507 | 5 | -3 | 13 | 0.22 |
| | 26 | 20% | 5402 | 5 | -5 | 14 | 0.33 |
| | 26 | 40% | 5569 | 8 | -3 | 19 | 0.16 |
| | 26 | 60% | 4977 | 0 | -11 | 11 | 0.96 |
| | 52 | 0% | 5316 | 3 | -5 | 12 | 0.44 |

| | | | | | | | |
|---|---|---|---|---|---|---|---|
| | 52 | 20% | 4840 | -2 | -10 | 8 | 0.7 |
| | 52 | 40% | 5180 | 2 | -7 | 11 | 0.66 |
| | 52 | 60% | 5097 | 1 | -10 | 13 | 0.81 |
| 26 | 26 | 0% | 5381 | 5 | -5 | 14 | 0.35 |
| | 26 | 20% | 4919 | -1 | -11 | 8 | 0.84 |
| | 26 | 40% | 5303 | 4 | -7 | 15 | 0.46 |
| | 26 | 60% | 4990 | 0 | -12 | 13 | 0.98 |
| | 52 | 0% | 4575 | -4 | -13 | 4 | 0.3 |
| | 52 | 20% | 4916 | -1 | -11 | 9 | 0.84 |
| | 52 | 40% | 5157 | 2 | -8 | 11 | 0.7 |
| | 52 | 60% | 4747 | -3 | -16 | 9 | 0.54 |
| 52 | 26 | 0% | 4968 | 0 | -10 | 10 | 0.94 |
| | 26 | 20% | 4577 | -6 | -16 | 5 | 0.3 |
| | 26 | 40% | 4976 | 0 | -13 | 14 | 0.95 |
| | 26 | 60% | 4655 | -6 | -20 | 9 | 0.4 |
| | 52 | 0% | 4302 | -7 | -15 | 1 | 0.09 |
| | 52 | 20% | 4234 | -8 | -17 | 0 | 0.06 |
| | 52 | 40% | 4383 | -7 | -15 | 2 | 0.13 |
| | 52 | 60% | 4128 | -11 | -21 | -1 | **0.03** |

*Note: All tests are comparing the given scenario for reactive underpass closures (table rows) against the baseline scenario. Statistically significant results are highlighted by bold text for* p-*values. Color indicates the direction of change: red indicates that the given reactive underpass closure scenario significantly increased epidemic duration; blue that closures significantly reduced epidemic duration. The duration of closure column indicates the duration (in weeks) that simulated underpasses were closed. Onset of reactive is the time (in weeks) after epidemic* *initiation at which reactive closures began; the proportion proactive column gives the proportion of population proactively vaccinated prior to an epidemic. MW test statistic is the Mann-Whitney* U *test statistic. The Estimate column gives the Mann-Whitney* U *difference estimate, and the 95% confidence interval is given in the 95% CI (L) (lower) and 95% CI (U) (upper) columns.*

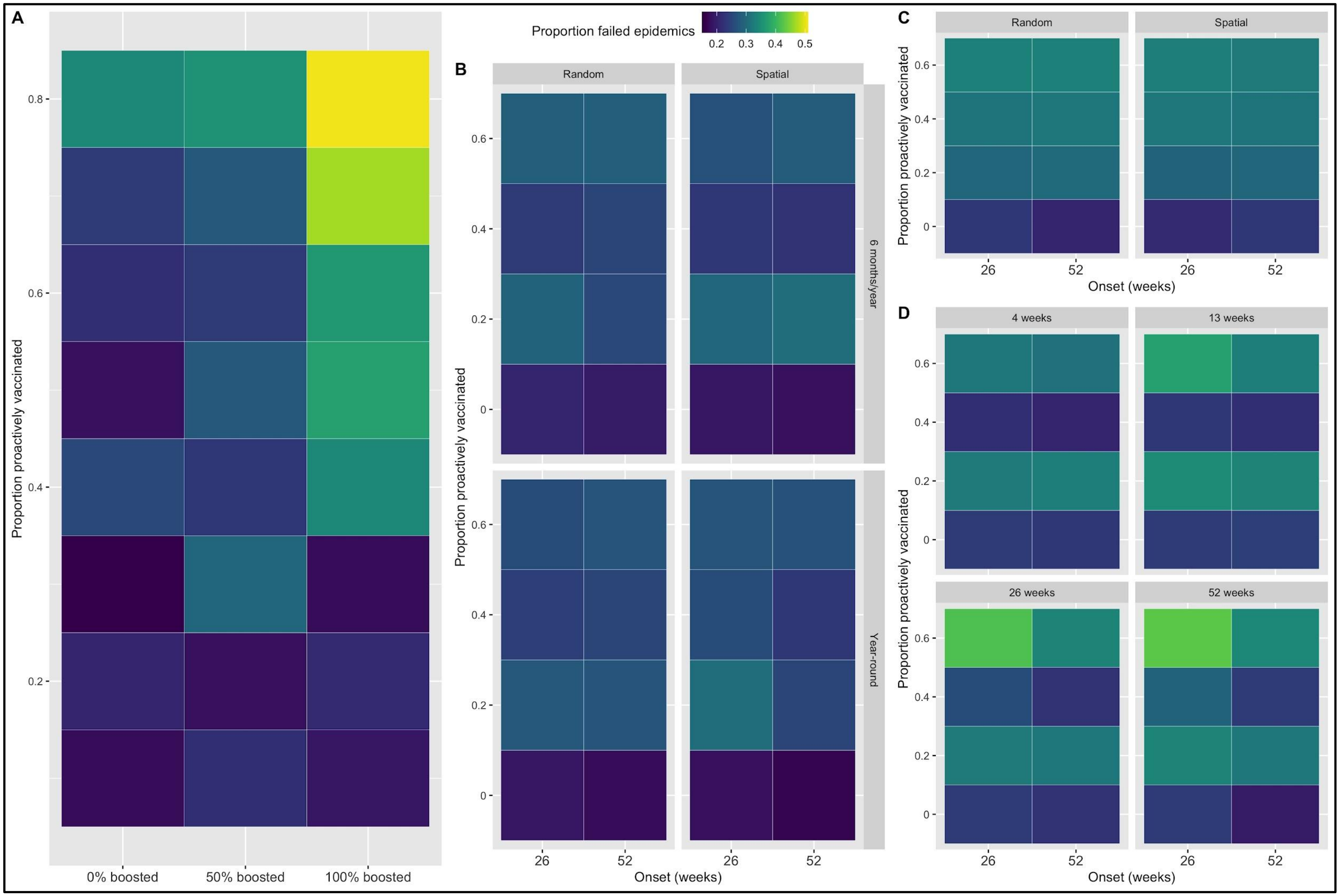

**Figure S17:** Heat maps of the proportion of simulated FeLV epidemics that failed (fewer than 5 progressive or regressive infections) per 100 successful epidemics. Results are shown for (A) proactive vaccination alone; (B) reactive and proactive vaccination; (C)

reactive test-and-removal with proactive vaccination; and (D) reactive underpass closures with proactive vaccination. In all plots, the y-axis gives the proportion of the population proactively vaccinated. For (A), the x-axis gives the proportion of individuals receiving boosted vaccination; in all other plots, 50% of all vaccinated individuals received boosted vaccinations. For plots B-D, the x-axis gives the onset of reactive interventions after epidemic initiation (26 or 52 weeks). In B and C, panel columns represent strategy for reactive vaccination and captures/testing, respectively. In addition, panel rows in B show the duration of reactive vaccination per year. Lastly, each panel in D represents the duration of underpass closures. Each colored square represents the results of 100 simulations.

*Sensitivity analysis of no-intervention scenarios*

As reported in the main text, simulated FeLV outbreak sizes were variable across the 50 sensitivity analysis parameter sets examined under a no-intervention scenario (Figure S18). PRCC results identified network density (Net_dens), infectiousness of regressives (*C*), weekly contact rates ($\omega$), and the baseline probability of transmission ($\beta$) as statistically significant parameters with positive correlations to median mortalities; the weekly probability of mortality ($\mu$) showed a significant negative correlation with median mortalities (Figure S19). Note, however, that PRCC inference may be affected by discrete parameters (Marino et al., 2008), so these results should be interpreted with some caution.

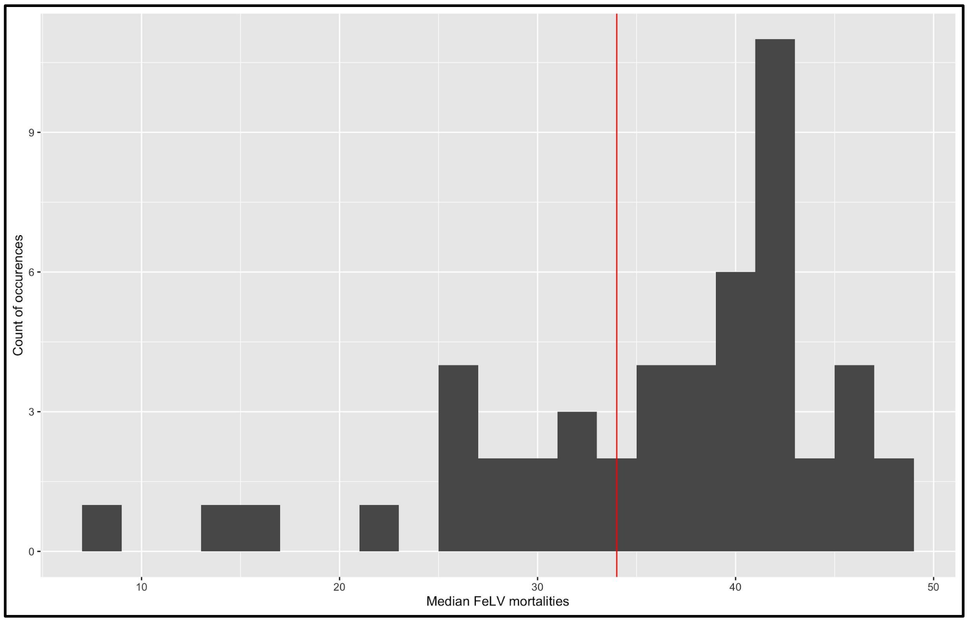

**Figure S18:** Histogram of median simulated FeLV mortalities for each of 50 sensitivity analysis parameter sets evaluated under a no-intervention scenario. The vertical red line represents the median mortalities with no interventions from our main analysis simulations.

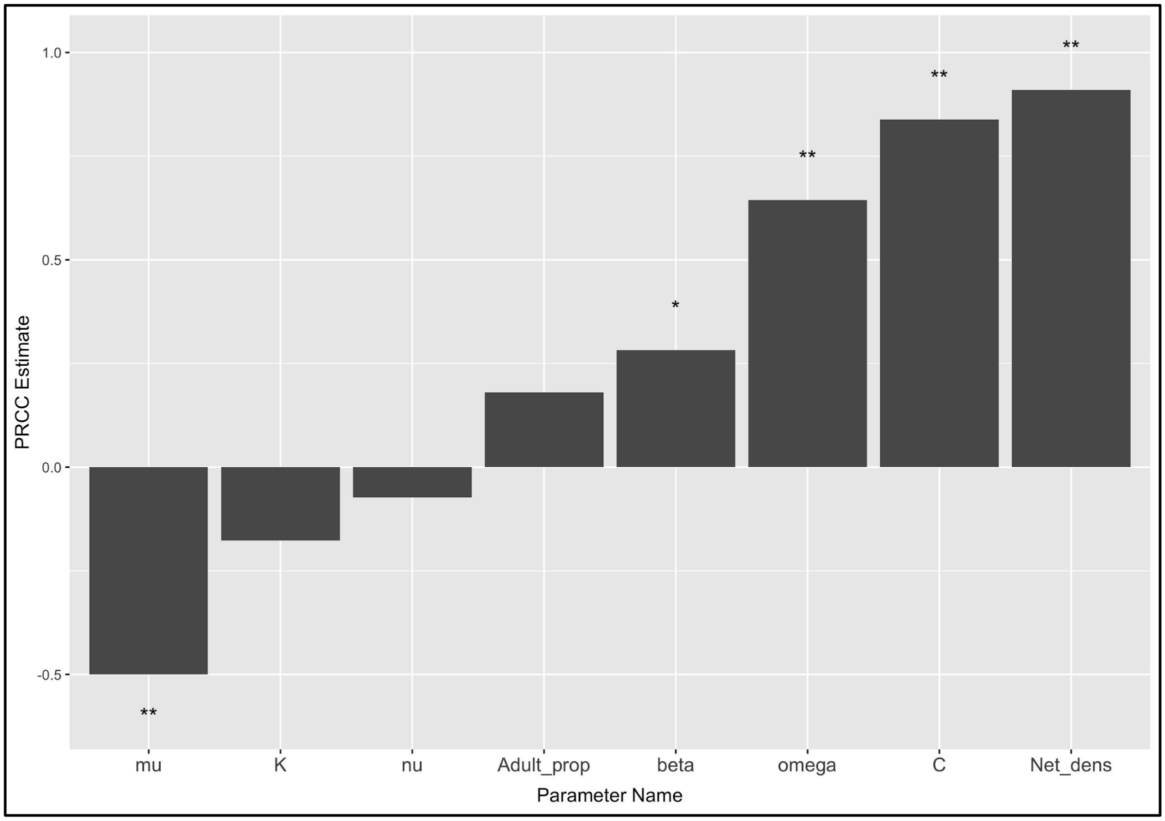

**Figure S19:** Partial Rank Correlation Coefficient (PRCC) estimates for simulation parameters in no-intervention scenario sensitivity analysis. The outcome considered was the median mortalities per parameter set. Parameter names and descriptions can be found in Table S1. Asterisks indicate statistical significance: * = $p \leq 0.05$; ** = $p \leq 0.01$.

*Post-hoc sensitivity analysis*

In our initial sensitivity analysis of proactive vaccination scenarios, we found that our results varied qualitatively from our main simulations, with low levels of proactive vaccination sometimes effective at reducing FeLV outbreak sizes. Focusing on the subset of sensitivity scenarios in which 20% of the population was proactively vaccinated with 100% boosting, we examined scatterplots and PRCC for parameter importance in relation to the outcome of the difference between median mortalities with and without proactive vaccination (hereafter, *mortality difference*; i.e., a difference value greater than 0 means that 20% vaccination with

100% boosting increased mortalities). PRCC analysis identified parameters of network density, infectiousness for regressives, and weekly contact rates (Net_dens, *C*, and $\omega$, respectively) as statistically significant parameters positively correlated with mortality difference at 20% proactive vaccination with 100% boosting (Figure S20). However, PRCC inference may be affected by discrete parameters and may be confounded by non-monotonic data (Marino et al., 2008). We therefore also evaluated scatterplots of our outcome across parameter values, finding that the relationship between mortality difference and network density, in particular, was not obviously monotonic, though data was limited (Figure S21). Given these limitations, we completed additional simulations to further evaluate the effect of network density on the mortality difference outcome.

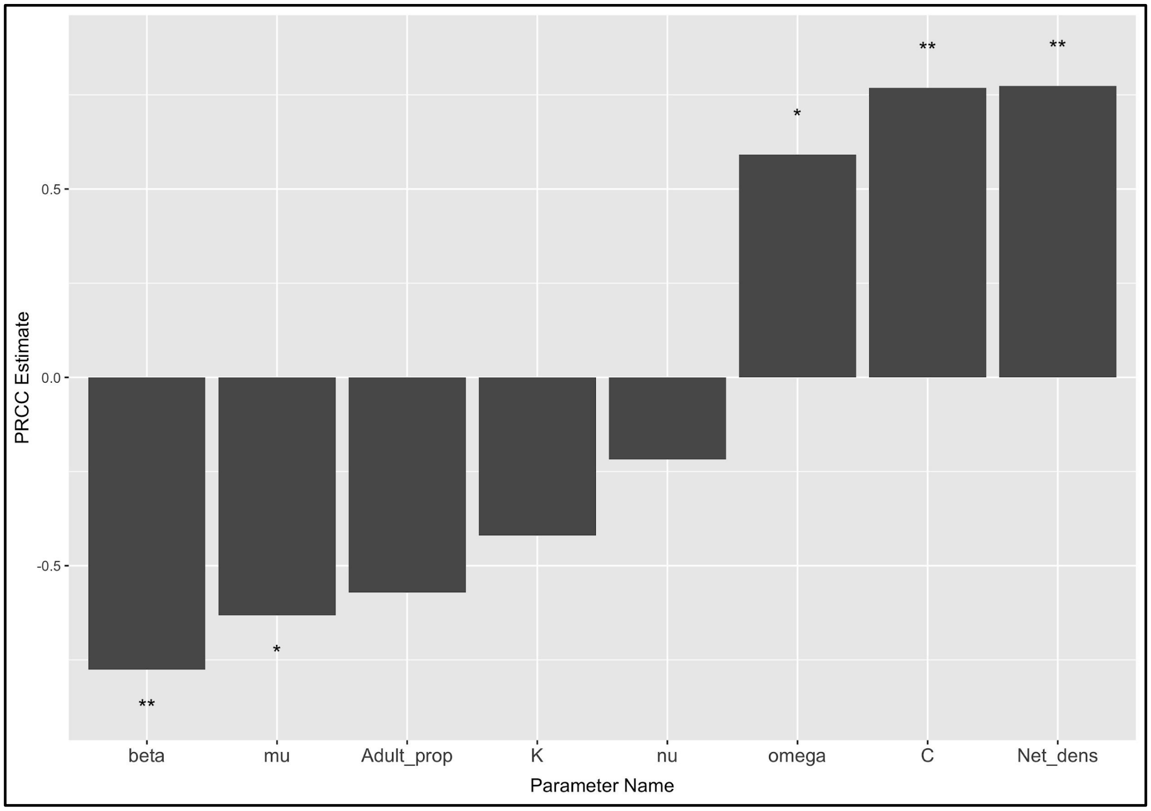

**Figure S20:** Partial Rank Correlation Coefficient (PRCC) estimates for simulation parameters in proactive vaccination scenario sensitivity analysis. The outcome considered was the difference in median mortalities with and without proactive vaccination (per parameter set). Shown are results for parameter sets in which 20% of the population was proactively vaccinated with 100% boosting. Parameter names and descriptions can be found in Table S1. Asterisks indicate statistical significance: * = $p \le 0.05$; ** = $p \le 0.01$.

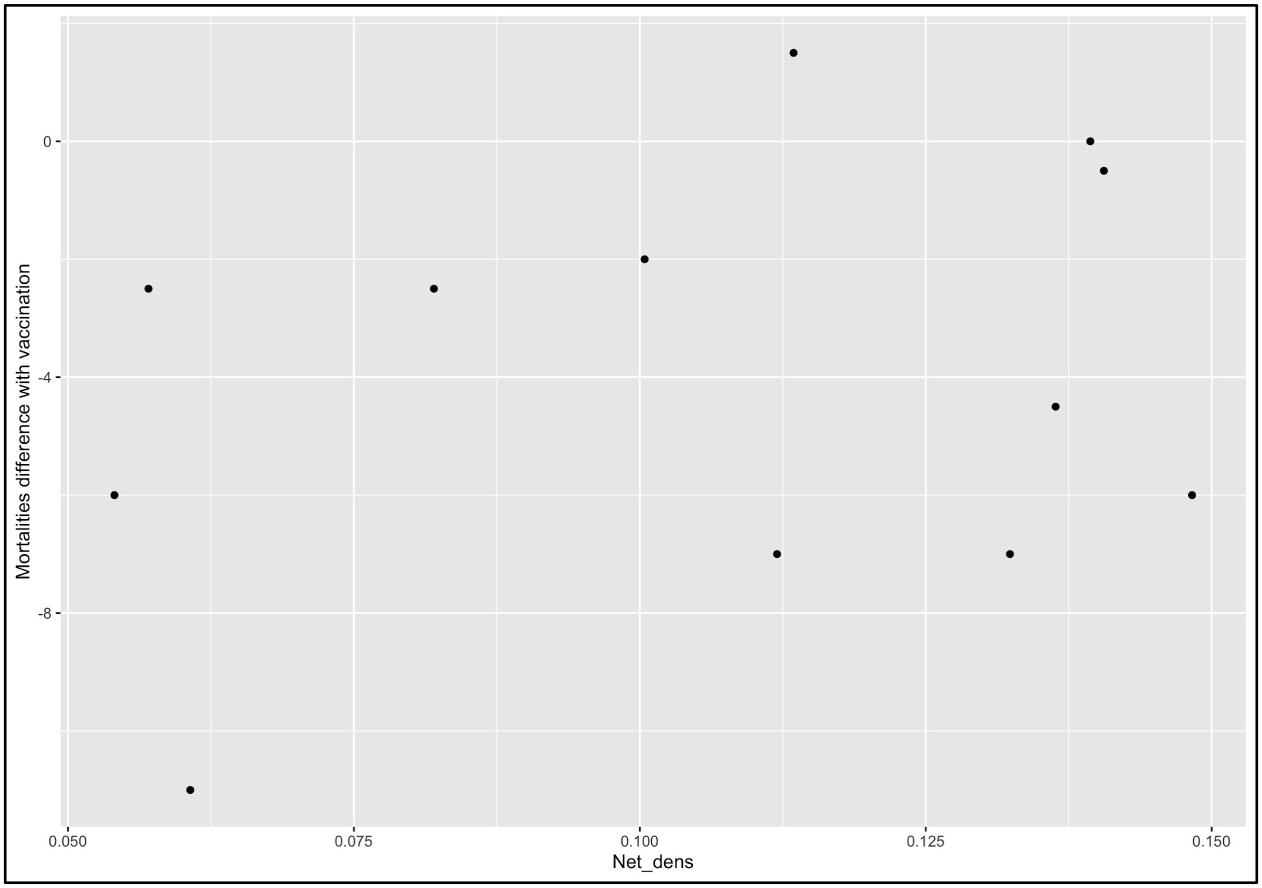

**Figure S21:** Scatterplot of difference in median mortalities with and without proactive vaccination (“mortalities difference” per parameter set) against network density parameterization. Shown are results for parameter sets in which 20% of the population was proactively vaccinated with 100% boosting.

To determine parameter space to target for additional simulations—and potential additional non-monotonicities—we first classified the results for each proactive vaccination sensitivity analysis parameter set (n=12) according to if low levels of proactive vaccination qualitatively reduced mortalities, made little to no change, or increased mortalities. We then examined these classifications across the parameter space represented by each pair of these 12 parameter sets (Figure S22) to determine if any parameters or pairs of parameters showed clustering of classifications in parameter space. We were limited to this qualitative analysis by computational complexity (these 12 parameter sets alone required 5,400 simulations), but found some indications that, in addition to the previously identified target parameter of network density, the parameter for proportion adults also appeared to be associated with classification of simulation results as either “little to no change in mortalities” or “increased mortalities” at low levels of vaccination (Figure S22).

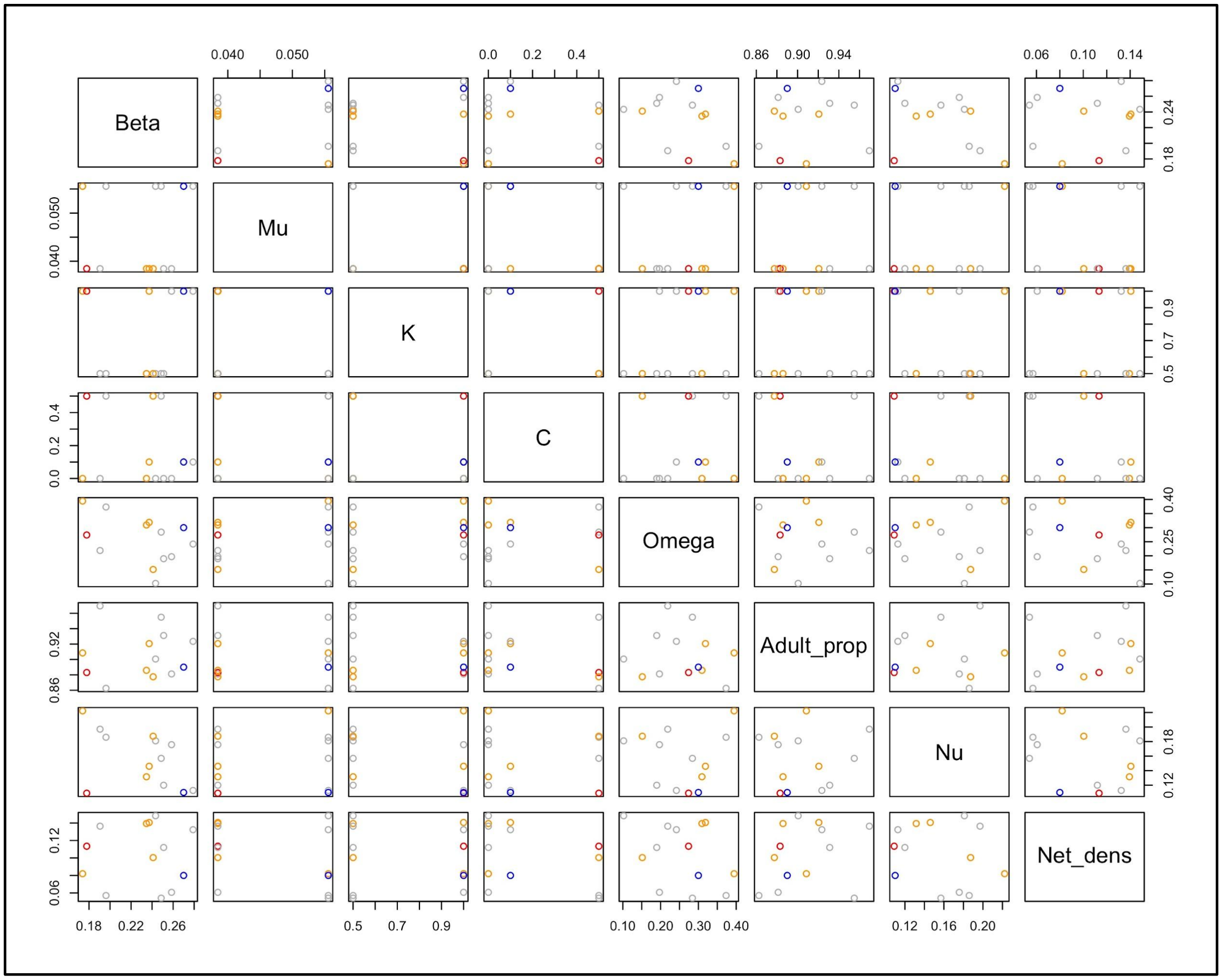

**Figure S22:** Plot of proactive vaccination sensitivity analysis parameter set classifications across all pairs of parameters. Points represent individual parameter sets for the main sensitivity analysis of proactive vaccination (n = 12), and colors represent qualitative classifications of results. The blue point per plot shows the parameter space represented by our main simulations. For all other points, red = increased mortalities with low levels of proactive vaccination; orange = little to no change in mortalities with low levels of proactive vaccination; gray = reduced mortalities with low levels of proactive vaccination.

To further examine this potential relationship, we generated an additional two parameter sets for a *post-hoc* sensitivity analysis of proactive vaccination scenarios (Table S2). In the first set, we used the same target parameters from our main analyses (Table S1) for all parameters except network density and proportion adults. The latter two parameters were assigned based on apparent intermediate parameter space, which we hypothesized should preferentially result in low levels of proactive vaccination being associated with increased mortalities. This resulted in an additional 4 parameter sets across a no-intervention baseline and 9 variations in proactive vaccination (20/40/60% of the population vaccinated with 0/50/100% of individuals receiving boosted inoculations); with 50 simulations for each variation, this totaled an additional 2,000 simulations.

The second additional sensitivity parameter set held network density and proportion adults parameters constant from our main analyses (Table S1), but drew all other simulation parameters from the seven proactive vaccination sensitivity analysis parameter sets that qualitatively produced results most divergent from our main analyses (Table S2). Here, we hypothesized that if network density and proportion adults really are important drivers behind our results, these simulations should now produce results more qualitatively aligned with our main analyses. This resulted in an additional 7 parameter sets across a no-intervention baseline and 6 variations in proactive vaccination (20/40/60% of the population vaccinated with 50/100% of individuals receiving boosted inoculations; we did not include 0% boosted in order to reduce computational intensity). With 50 simulations for each variation, this totaled an additional 2,450 simulations.

For both *post-hoc* sensitivity analysis parameter sets, we examined qualitative alignment with our main results to determine if particular parameter space was associated with our main finding that low levels of proactive vaccination could paradoxically increase FeLV outbreak mortalities. These additional analyses found some evidence that intermediate values for network structural parameters (network density and proportion adults) may be associated with the qualitative result that low proactive vaccination can increase FeLV mortalities; this may be especially true under conditions of high transmission potential (e.g. higher infectiousness of regressive individuals and/or increased weekly contact rates; Figure S23). While our *post-hoc* sensitivity analysis did not specifically examine individual transmission parameters, we do note that high weekly probabilities of contact are also tentatively associated with our qualitative results (Figure S24), which is consistent with our initial PRCC analysis with proactive vaccination sensitivity scenarios.

**Table S10: Post-Hoc Sensitivity Analysis Parameter Sets**

| | Set Name | *Post-Hoc* Set 1 | | | | *Post-Hoc* Set 2 | | | | | | |
|---|---|---|---|---|---|---|---|---|---|---|---|---|
| | **Set Number** | 1 | 2 | 3 | 4 | 1 | 2 | 3 | 4 | 5 | 6 | 7 |
| **Parameter Name** | Adult_prop | 0.8875 | 0.8875 | 0.9125 | 0.9125 | 0.89 | 0.89 | 0.89 | 0.89 | 0.89 | 0.89 | 0.89 |
| | Net_dens | 0.0875 | 0.1125 | 0.0875 | 0.1125 | 0.08 | 0.08 | 0.08 | 0.08 | 0.08 | 0.08 | 0.08 |
| | $\beta$ | 0.27 | 0.27 | 0.27 | 0.27 | 0.259 | 0.243 | 0.249 | 0.196 | 0.251 | 0.191 | 0.279 |
| | C | 0.1 | 0.1 | 0.1 | 0.1 | 0 | 0 | 0.5 | 0.5 | 0 | 0 | 0.1 |
| | $\omega$ | 0.3 | 0.3 | 0.3 | 0.3 | 0.197 | 0.102 | 0.284 | 0.373 | 0.190 | 0.219 | 0.242 |
| | $\mu$ | 1/18 | 1/18 | 1/18 | 1/18 | 1/26 | 1/18 | 1/18 | 1/18 | 1/26 | 1/26 | 1/18 |
| | K | 1 | 1 | 1 | 1 | 1 | 0.5 | 0.5 | 0.5 | 0.5 | 0.5 | 1 |
| | $\nu$ | 0.11 | 0.11 | 0.11 | 0.11 | 0.176 | 0.181 | 0.167 | 0.186 | 0.120 | 0.197 | 0.113 |
| | P | 0.25 | 0.25 | 0.25 | 0.25 | 0.25 | 0.25 | 0.25 | 0.25 | 0.25 | 0.25 | 0.25 |

*Note: Parameter definitions can be found in Table S1.*

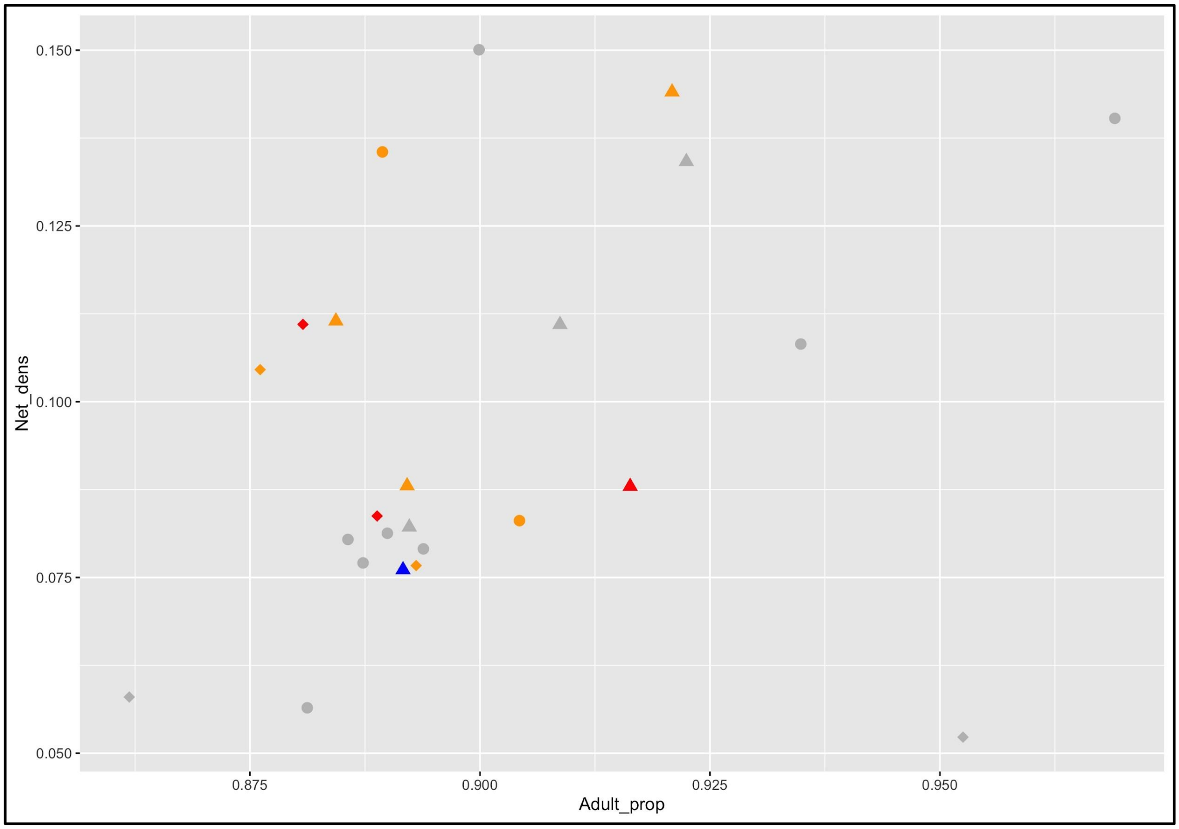

**Figure S23:** Scatterplot of *post-hoc* sensitivity analysis evaluation of influence of proportion adults (Adult_prop) and network density (Net_dens) parameters. Points represent individual parameter sets for both the main sensitivity analysis of proactive vaccination (n=12) and both *post-hoc* parameter sets (n = 4 and 7). Point colors represent qualitative classifications of results. Blue shows the parameter space represented by our main simulations. For all other points, red = increased mortalities with low levels of proactive vaccination; orange = little to no change in mortalities with low levels of proactive vaccination; gray = reduced mortalities with low levels of proactive vaccination. Point shapes give the value for the constant modifying infectiousness of regressives (parameter C): circle = 0, triangle = 0.1, diamond = 0.5. Note: points are jittered for visibility.

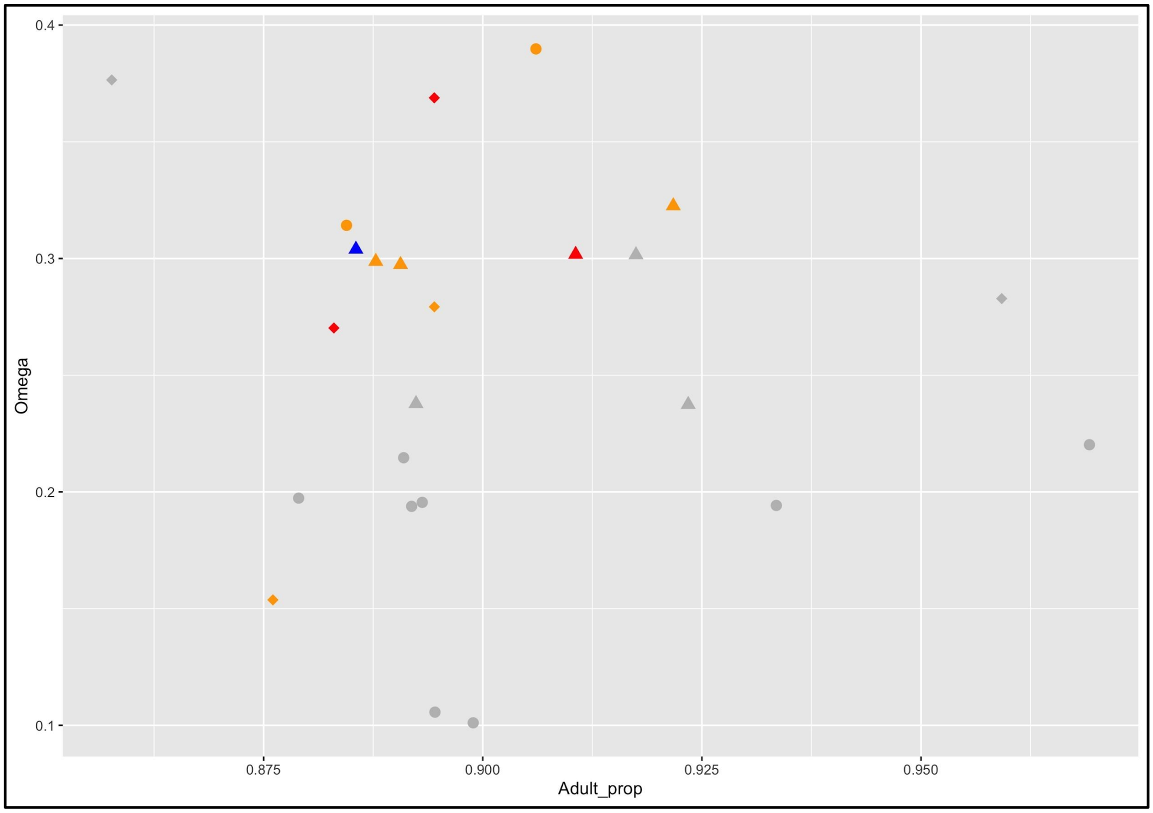

**Figure S24:** Scatterplot of *post-hoc* sensitivity analysis results, here showing the proportion adults (Adult_prop) against the weekly probability of contact (Omega/$\omega$) parameters, though $\omega$ was not specifically evaluated in *post-hoc* analysis. Points represent individual parameter sets for both the main sensitivity analysis of proactive vaccination (n=12) and both *post-hoc* parameter sets (n = 4 and 7). Point colors represent qualitative classifications of results. Blue shows the parameter space represented by our main simulations. For all other points, red = increased mortalities with low levels of proactive vaccination; orange = little to no change in mortalities with low levels of proactive vaccination; gray = reduced mortalities with low levels of proactive vaccination. Point shapes give the value for the constant modifying infectiousness of regressives (parameter C): circle = 0, triangle = 0.1, diamond = 0.5. Note: points are jittered for visibility.

### Discussion

*FeLV transmission*

While our sensitivity analysis highlighted the impacts of network structure on pathogen management requirements, additional uncertainty in FeLV transmission parameters should be considered. In particular, our main simulations allowed limited transmission from regressively infected individuals, which contrasts with expectations from domestic cats, but was supported by model-based evidence in Gilbertson *et al.* (2021). In addition, we assumed only a single spillover event from domestic cats, but these events occur with uncertain frequency (Chiu et al., 2019). The virulence of any circulating FeLV strain may also affect model predictions. There was some evidence that the FeLV strain in the 2002-2004 outbreak among panthers was particularly virulent (Brown et al., 2008). However, the same does not appear to have been true for Iberian lynx (Geret et al., 2011). Severity of FeLV-induced disease may therefore reflect the effects of inherent genetic susceptibility, particularly in these highly inbred populations (Geret et al., 2011; Johnson et al., 2010; Roelke et al., 1993). Lower virulence FeLV strains than we have modeled here and/or increased population genetic diversity may therefore reduce severity of disease and even transmission potential in panthers.

*Temporary spatial restrictions are unlikely to be effective under realistic scenarios*

Here, we examined the effect of wildlife underpass closures as a novel method to restrict connectivity of the panther population under emergency disease control conditions. Such temporary spatial restrictions could increase other types of panther mortality (e.g. vehicle strikes or intraspecific conflict), so such an intervention would need to reduce FeLV mortalities at a rate greater than these other types of panther mortality in order to be a viable disease control strategy. Unfortunately, our simulations found that temporary underpass closures were generally no more effective at reducing FeLV mortalities than less risky reactive interventions. Further, underpass closures were less effective when occurring after the peak of simulated epidemics, such that spatial restrictions would need to occur early in an epidemic to be effective. However, it should be noted that spatial restrictions may be more effective if used in combination with other reactive FeLV management strategies (e.g., reactive vaccination in concert with temporary spatial restrictions to prevent transmission resurgence after restrictions are removed).

Extensive research has considered how landscape barriers and fragmentation may affect pathogen transmission and control in wildlife (e.g., Rees et al., 2013; Smith et al., 2002; Tracey et al., 2014; White et al., 2018). By closing underpasses, we artificially fragmented panther habitat, assuming complete efficacy of closures in preventing transmission across the I-75 freeway. In reality, some individuals would likely successfully traverse this barrier. Because habitat fragmentation can promote pathogen persistence (White et al., 2018), a "semipermeable" freeway barrier may, in fact, ultimately worsen epidemic dynamics. Alternatively, landscape restrictions to animal movement can facilitate more efficient disease control, as in Haydon *et al.* (2006), where vaccinating to control pathogen spread along habitat corridors reduced the level of vaccine coverage needed to protect endangered Ethiopian wolves. While panthers are generally well-connected spatially, expansion of the population north of its current range may favor similar metapopulation-oriented disease control strategies. However, given uncertainties in the effect of underpass closures on other sources of mortalities and their limited effectiveness in our simulations, we suggest temporary underpass closures should currently be considered a low priority for FeLV management in panthers.

### Supplementary References

Brown, M. A., Cunningham, M. W., Roca, A. L., Troyer, J. L., Johnson, W. E., & O'Brien, S. J. (2008). Genetic characterization of feline leukemia virus from Florida panthers. *Emerging Infectious Diseases*, *14*(2), 252–259.

Chiu, E. S., Kraberger, S., Cunningham, M., Cusack, L., Roelke, M., & VandeWoude, S. (2019). Multiple Introductions of Domestic Cat Feline Leukemia Virus in Endangered Florida Panthers. *Emerging Infectious Diseases*, *25*(1), 92–101.

Cotterill, G. G., Cross, P. C., Cole, E. K., Fuda, R. K., Rogerson, J. D., Scurlock, B. M., & du Toit, J. T. (2018). Winter feeding of elk in the Greater Yellowstone Ecosystem and its effects on disease dynamics. *Philosophical Transactions of the Royal Society of London. Series B, Biological Sciences*, *373*(1745). https://doi.org/10.1098/rstb.2017.0093

Geret, C.P., Cattori, V., Meli, M.L., Riond, B., Martinez, F., Lopez, G., Vargas, A., Simon, M.A., Lopez-Bao, J.V., Hofmann-Lehmann, R. and Lutz, H., (2011). Feline leukemia virus outbreak in the critically endangered Iberian lynx (*Lynx pardinus*): high-throughput sequencing of envelope variable region A and experimental transmission. *Archives of virology*, *156*(5), 839-854. https://doi.org/10.1007/s00705-011-0925-z

Gilbertson, M. L. J., Fountain-Jones, N. M., Malmberg, J. L., Gagne, R. B., Lee, J. S., Kraberger, S., Kechejian, S., Petch, R., Chiu, E., Onorato, D., Cunningham, M. W., Crooks, K. R., Funk, W. C., Carver, S., VandeWoude, S., VanderWaal, K., & Craft, M. E. (2021). Transmission of one predicts another: Apathogenic proxies for transmission dynamics of a fatal virus. *bioRxiv*. https://doi.org/10.1101/2021.01.09.426055

Haydon, D. T., Randall, D. A., Matthews, L., Knobel, D. L., Tallents, L. A., Gravenor, M. B., Williams, S. D., Pollinger, J. P., Cleaveland, S., Woolhouse, M. E. J., Sillero-Zubiri, C., Marino, J., Macdonald, D. W., & Laurenson, M. K. (2006). Low-coverage vaccination strategies for the conservation of endangered species. *Nature*, *443*(7112), 692–695.

Johnson, W. E., Onorato, D. P., Roelke, M. E., Land, E. D., Cunningham, M., Belden, R. C., McBride, R., Jansen, D., Lotz, M., Shindle, D., Howard, J., Wildt, D. E., Penfold, L. M., Hostetler, J. A., Oli, M. K., & O'Brien, S. J. (2010). Genetic restoration of the Florida panther. *Science*, *329*(5999), 1641–1645.

Keeling, M. J., & Eames, K. T. D. (2005). Networks and epidemic models. *Journal of the Royal Society, Interface / the Royal Society*, *2*(4), 295–307.

Lloyd-Smith, J. O., Schreiber, S. J., Kopp, P. E., & Getz, W. M. (2005). Superspreading and the effect of individual variation on disease emergence. *Nature*, *438*(7066), 355–359.

Marino, S., Hogue, I. B., Ray, C. J., & Kirschner, D. E. (2008). A methodology for performing global uncertainty and sensitivity analysis in systems biology. *Journal of Theoretical Biology*, *254*(1), 178–196.

Mysterud, A., & Rolandsen, C. M. (2019). Fencing for wildlife disease control. *The Journal of Applied Ecology*, *56*(3), 519–525.

Rees, E. E., Pond, B. A., Tinline, R. R., & Bélanger, D. (2013). Modelling the effect of landscape heterogeneity on the efficacy of vaccination for wildlife infectious disease control. *The Journal of Applied Ecology*, *50*(4), 881–891.

Roelke, M. E., Forrester, D. J., Jacobson, E. R., Kollias, G. V., Scott, F. W., Barr, M. C., Evermann, J. F., & Pirtle, E. C. (1993). Seroprevalence of infectious disease agents in free-ranging Florida panthers (*Felis concolor coryi*). *Journal of Wildlife Diseases*, *29*(1), 36–49.

Smith, D. L., Lucey, B., Waller, L. A., Childs, J. E., & Real, L. A. (2002). Predicting the spatial dynamics of rabies epidemics on heterogeneous landscapes. *Proceedings of the National Academy of Sciences of the United States of America*, *99*(6), 3668–3672.

Tracey, J. A., Bevins, S. N., VandeWoude, S., & Crooks, K. R. (2014). An agent-based movement model to assess the impact of landscape fragmentation on disease transmission. *Ecosphere*, *5*(9), 1–24.

White, L. A., Forester, J. D., & Craft, M. E. (2018). Disease outbreak thresholds emerge from interactions between movement behavior, landscape structure, and epidemiology. *Proceedings of the National Academy of Sciences of the United States of America*, *115*(28), 7374–7379.

Wu, J., Dhingra, R., Gambhir, M., & Remais, J. V. (2013). Sensitivity analysis of infectious disease models: methods, advances and their application. *Journal of the Royal Society, Interface / the Royal Society*, *10*(86), 20121018.
